# Y705 and S727 are required for mitochondrial import and transcriptional activities of STAT3 and regulate proliferation of embryonic and tissue stem cells

**DOI:** 10.1101/2020.07.17.208264

**Authors:** Margherita Peron, Giacomo Meneghetti, Alberto Dinarello, Laura Martorano, Riccardo M. Betto, Nicola Facchinello, Annachiara Tesoriere, Natascia Tiso, Graziano Martello, Francesco Argenton

## Abstract

The STAT3 transcription factor, acting both in the nucleus and mitochondria, maintains embryonic stem cell pluripotency and promotes their proliferation. In this work, using zebrafish, we determined *in vivo* that mitochondrial STAT3 regulates mtDNA transcription in embryonic and larval stem cell niches and that this activity affects their proliferation rates. As a result, we demonstrated that STAT3 import inside mitochondria requires Y705 phosphorylation by Jak2, while its mitochondrial transcriptional activity, as well as its effect on proliferation, depends on the MAPK target S727. These data were confirmed using mouse embryonic stem cells: Y705 mutated STAT3 cannot enter the mitochondrion while the S727 mutation does not affect mitochondrial import of the protein. Surprisingly, STAT3-dependent increase of mitochondrial transcription seems independent from STAT3 binding to STAT3 responsive elements. Finally, loss of function experiments, with chemical inhibition of JAK/STAT3 pathway or genetic ablation of *stat3* gene, demonstrated that STAT3 is also required for cell proliferation in the intestine of zebrafish.

## INTRODUCTION

The investigation of the role of Signal Transducer and Activator of Transcription 3 (STAT3) pathway in human diseases represents, to date, one of the most exciting developments in modern medicine (O’Shea *et al.*, 2015). Many of the major human malignancies display elevated levels of constitutively activated STAT3 (Johnston and Grandis, 2011). Most interestingly, recent data report that STAT3 target genes are overexpressed in tumour-initiating cancer stem cells (Fouse and Costello, 2013; Wei *et al.*, 2014; Ghoshal *et al.*, 2016). Stat3 is also the key mediator of Leukemia Inhibitory Factor (LIF) in mouse embryonic stem cells (ESCs) where the LIF/STAT3 axis promotes the maintenance and induction of naïve pluripotency (Burdon *et al.*, 1999; Matsuda *et al.*, 1999; Smith *et al.*, 1988; Martello *et al.*, 2013).

STAT3 transcriptional activity is regulated by phosphorylation of two separate residues. When Janus kinases 1/2/3 (JAK1/2/3) phosphorylate its tyrosine 705 (Y705), STAT3 dimerizes, enters the nucleus, binds to STAT3 response elements and triggers transcription of its target genes (Ni *et al.*, 2004). On the other hand, the function of serine 727 (S727) phosphorylation remains controversial; pS727 has been reported to have both activating and inhibitory effects on STAT3 transcriptional activity (Quin *et al.*, 2008; Shi *et al.*, 2006). More recently, it has been demonstrated that pY705 is absolutely required for STAT3-mediated ESCs self-renewal, while pS727 is dispensable, serving only to promote proliferation and optimal pluripotency (Huang *et al.*, 2014). Notably, zebrafish mutants lacking maternal and zygotic Stat3 expression display transient axis elongation defects due to reduced cell proliferation during embryogenesis (Liu *et al.*, 2017). Additionally, it has been demonstrated that in zebrafish Stat3 is transcriptionally active in stem cells of highly proliferative tissues like Tectum Opticum (TeO), hematopoietic tissue and intestine (Peron *et al.*, 2020).

Besides its canonical nuclear functions, a pool of STAT3 has been reported also in the mitochondrion of different cell types, thus including this transcription factor in the large family of dual-targeted proteins with both nuclear and mitochondrial functions (Wegrzyn *et al.*, 2009; Mantel *et al.*, 2012). A recently discovered subcellular localization of STAT3 is the Endoplasmic Reticulum (ER), where it controls the release of Ca^2+^, with consequences on the mitochondrial Ca^2+^ levels and on the life-death cell decision. This function is crucial for the maintenance in the tumour niche of apoptosis-resistant cells (Avalle *et al.*, 2019). Although the mechanisms of action of mitochondrial STAT3 (mitoSTAT3) are still debated, between 10 and 25% of total STAT3 has been shown to be in the mitochondria, (Szczepanek *et al.*, 2011; Carbognin et al., 2016). Using different *in vitro* models, several roles have been proposed for mitoSTAT3, such as: interaction with mitochondrial respiratory chain complexes I and II; binding to the d-loop regulatory region of mitochondrial DNA (mtDNA); regulation of mitochondrial gene expression; regulation of mitochondrial permeability transition pore (Wegrzyn *et al.*, 2009; Macias *et al.*, 2014; Meier and Larner, 2014; Carbognin *et al.*, 2016). However, the molecular mechanisms leading to mitoSTAT3 activation and translocation are still only partially understood. Previous *in vitro* studies suggested that the S727 phosphorylation by MAPK kinases may be required for STAT3 mitochondrial activity (Gough *et al.*, 2013) and it seems necessary to restore complexes I and II activities in *Stat3^-/-^* cells (Wegrzin *et al.*, 2009). Moreover, pS727 STAT3 targeted to the mitochondria is described to promote Ras-dependent transformation in human bladder carcinoma cells (Gough *et al.*, 2009). Notably, the role of other post-translational modifications on mitoSTAT3 activity have not been investigated yet.

Mostly due to its transparent body and external fertilization, Zebrafish is an organism widely used for analysis of gene expression and protein-gene functions. Knowing that the STAT3 protein and all its functional domains are highly conserved in zebrafish (Liang *et al.*, 2012, Oates *et al.*, 1999) we used this animal model to study the mitoSTAT3 pathway *in vivo.* In this paper we demonstrate the dependence of mitoSTAT3 activities from both Y705 and S727 phosphorylation, hence, that mitoSTAT3 mitochondrial function relies on both ERK and JAK1/2/3 kinases activation. Our data also show that mitoSTAT3 modulation of mitochondrial transcription does not require STAT3 DNA binding domain, consistently with the differences between the eukaryotic and the prokaryotic transcriptional machineries operating in the nucleus and mitochondria, respectively. Finally using zebrafish larvae, we directly linked the STAT3-dependent regulation of tissue stem cells proliferation to mitochondrial transcriptional activity.

## RESULTS

### mitoSTAT3 regulates cell proliferation in the PML of the TeO through mtDNA transcription

It is known that in keratinocytes and mouse ESCs a portion of STAT3 localizes to mitochondria, where it induces mitochondrial transcription and cell proliferation (Macias *et al.*, 2014; Carbognin *et al.*, 2016). Hence, we tested whether STAT3 and mitochondrial transcribed genes colocalize in proliferating regions of the zebrafish embryo, considering this colocalization a *“conditio sine qua non”* in a STAT3-mitochondrial-proliferation liaison. Facilitated by the fact that *mt_nd2* expression profile has already been described in zebrafish (Thisse *et al.*, 2001) and considering the polycistronic nature of mtDNA-transcribed genes, that results in stoichiometric mtDNA transcription (Taanman, 1999), we decided to use this mitochondrial gene as a hallmark of global mitochondrial gene expression. As described in Fig 1, *mt_nd2* and the cellular proliferation marker *pcna* are particularly expressed in regions also labelled by *Stat3* transcripts, such as the inner retina and the Peripheral Midbrain Layer (PML) of the Tectum Opticum (TeO) (Fig. 1; Fig. 2A), the progenitor source for tectal and torus neurons in the embryo (Galant *et al.*, 2016).

**Fig. 1:**
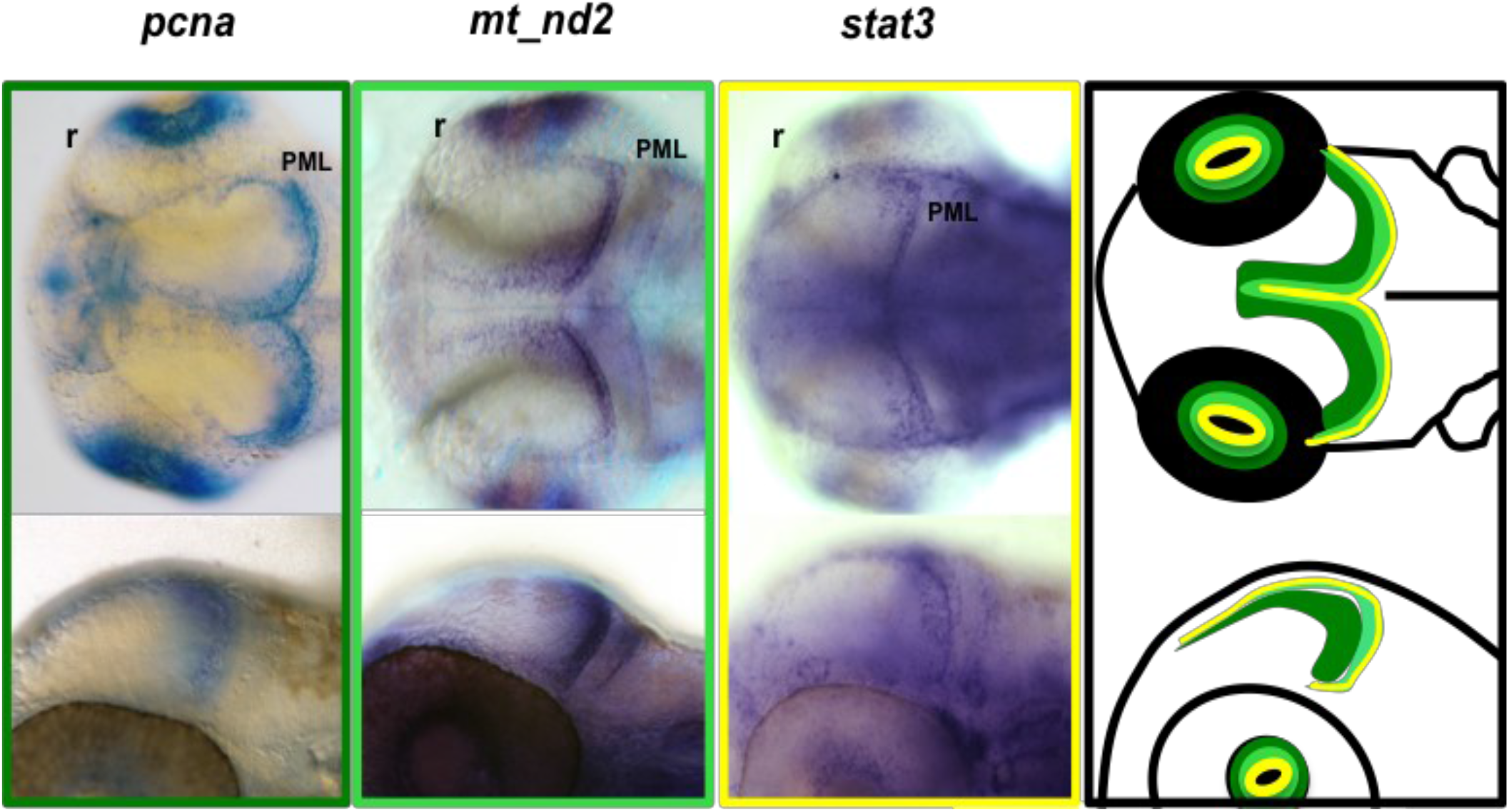
*Stat3* mRNA is co-expressed with proliferation and mtDNA transcription markers in the TeO of zebrafish embryos. Whole mount in situ hybridization (WISH) on 48-hpf zebrafish WT embryos using *pcna* (dark green frame and outline), *mt_nd2* (light green frame and outline), and *stat3* (yellow frame and outline) antisense mRNA probes shows co expression of the three transcripts in the PML region of the TeO; r=retina; PML= Peripheral Midbrain Layer.

**Fig. 2:**
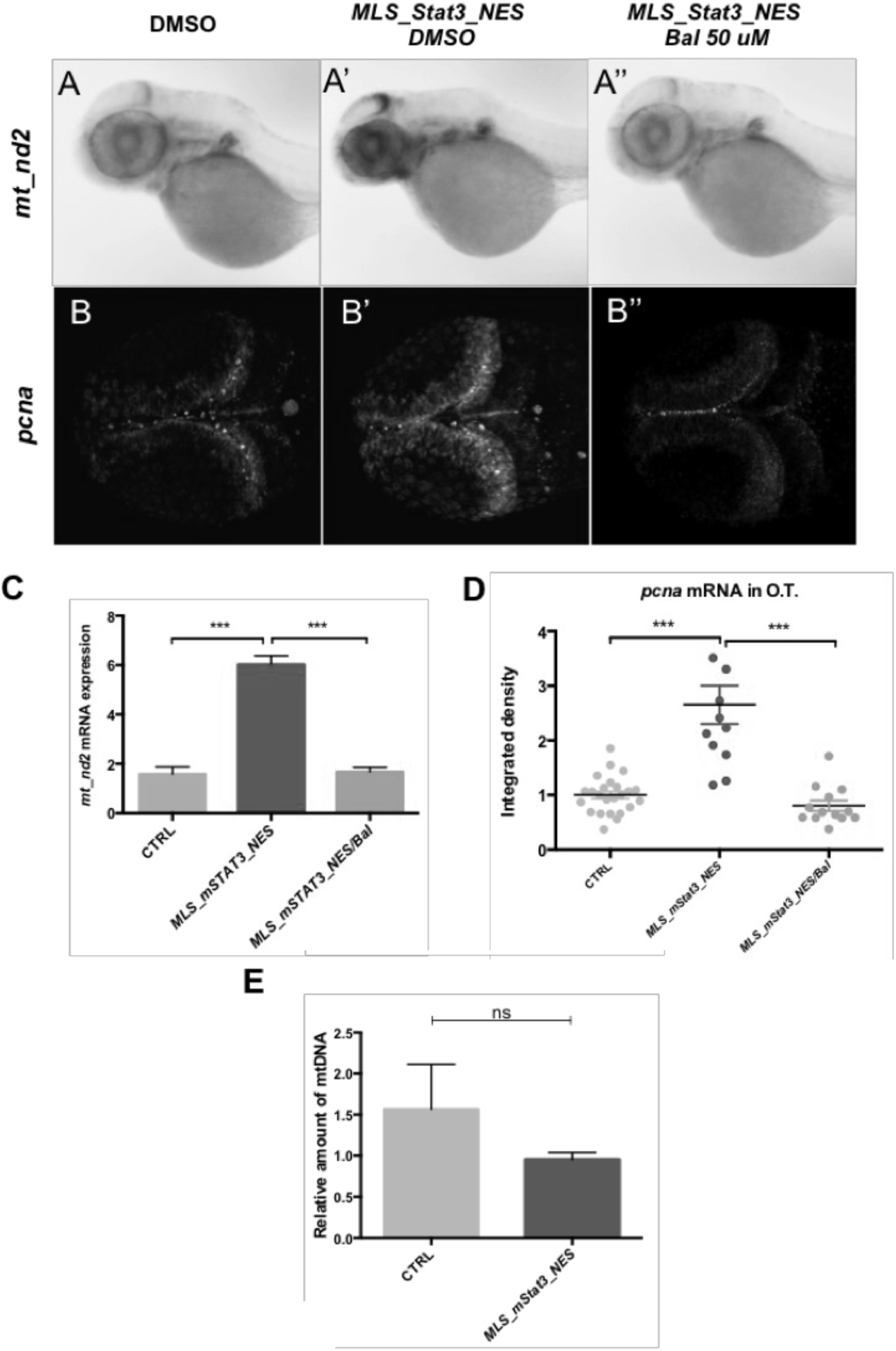
mitoSTAT3 regulates proliferation through mitochondrial DNA transcription. **A-A”:** WISH with anti-*mt_nd2* mRNA probe representing mitochondrial gene transcription in: uninjected embryos (A); embryos injected with *MLS_mStat3_NES* mRNA (A’); *MLS_mStat3_NES* mRNA injected embryos treated with 50 μM Balapiravir (A”). **B-B”:** Fluorescent *in situ* hybridization (FISH) with *pcna* probe in the TeO of: uninjected embryos (B); embryos injected with *MLS_mStat3_NES* mRNA (B’); and injected embryos treated with 50 μM Balapiravir (B”). **C:** qRT-PCR showing *mt_nd2* gene expression after injection of *MLS_mStat3_NES* mRNA and treatment with Balapiravir at 48 hours post injection (hpi); *zgapdh* was used as internal control (p-values= 0.0007; 0.0005). **D:** Fluorescence quantification of *pcna* mRNA expression in the TeO (n=12) (p-values= <0.0001; 0.0108; 0.0122). **E:** Relative amount of mtDNA in embryos injected with *MLS_mStat3_NES* mRNA and uninjected controls at 48 hpf. Mean dCt values were calculated as Ct of *mt_nd1* (mitochondrial encoded gene) minus Ct of *polg1* (nuclear encoded gene) (p-value= 0.3295). Statistical analysis was performed by unpaired t-test on 3 independent biological samples (where n not specified). ns: not significant; *p<0.05; **p<0.01; ***p<0.001; error bars=SEM.

In order to understand whether STAT3 mitochondrial activity regulates mtDNA transcription and promotes proliferation also *in vivo*, we injected zebrafish fertilized eggs with mRNA of a mitochondria-targeted murine form of *Stat3 (mStat3)*, provided with a nuclear export domain that makes it unable to localize to the nucleus *(MLS_mStat3_NES)* (Fig. S1 A). This chimeric protein a) is completely devoid of nuclear functions as assessed by qRT-PCR analysis of *Socs3*, the most direct *Stat3* target gene (Fig. S1 B) and b) efficiently localizes only inside the mitochondrion as revealed by its co-localization with the mitochondrial marker ATAD3, both in transfected mouse ESCs, and in zebrafish cells (Fig. S1 C; Fig. S2 A,B).

When *MLS_mStat3_NES* mRNA was injected into zebrafish embryos we could observe that this modified form of Stat3 was unable to induce the expression of its target gene *socs3a* (Fig. S3 A), however, we detect, both by *in situ* hybridization and qRT-PCR, a significant increase of mitochondrial transcription at 24 and 48 hours post fertilization (hpf) (Fig. 2A-A’, C; Fig. S3 B-D). It is worth noting that, as assayed by *pcna* analysis, overexpression of *MLS_mStat3_NES* mRNA also induced a proportional increase of proliferating cells in the same tissues where mitochondrial transcription was stimulated, i.e. the PML (Fig. 2 B-B’, C, D). On the other hand, we did not find any difference in mtDNA content when comparing injected and control embryos, suggesting that the effect of mitoSTAT3 on mitochondrial transcription is not due to increased mtDNA replication or mitochondrial biogenesis (Fig. 2E).

Importantly, chemical inhibition of mitochondrial transcription by using Balapiravir (Feng *et al.*, 2015) was able to abolish *MLS_Stat3_NES* effects on proliferation (Fig 2 A”, B”, D), thus providing evidence, *in vivo*, that replication of highly proliferating PML cells in the developing TeO of zebrafish embryos depends on mitoSTAT3-driven expression of mitochondrial genes.

### Mitochondrial STAT3 transcriptional activity relies on phosphorylation of both S727 and Y705

Additionally, we wanted to dissect the domains of STAT3 that are needed for the activation of mtDNA transcription, hence focussing our experiments on the DNA binding and the activation domain.

Putative STAT3 binding elements (SBE) have been identified in the mitochondrial D-Loop (the mitochondrial transcriptional initiation site) (Macias *et al.*, 2014) and STAT3 was found to immuno-precipitate together with the mitochondrial D-Loop in mouse ESCs (Carbognin *et al.*, 2016). Nonetheless, a form of mitochondrial STAT3 mutated in the DNA binding domain (STAT3 458-466 VVV-AAA) (MLS_mStat3_ΔDNAbd_NES), thus unable to bind SBE (Horvath *et al.*, 1995), retained its ability to activate *mt_nd2* transcription at comparable levels with respect to wild type (WT) mitoSTAT3 on zebrafish embryos (Fig. 3 A, A’). It is worth noting that this DNA binding mutant form when transfected in mouse ESCs is still able to co-localize with ATAD3, a mitochondrial nucleoid marker, (Fig. 3 B). This result suggests that binding of STAT3 to its specific response elements is dispensable for mtDNA transcription.

**Fig. 3:**
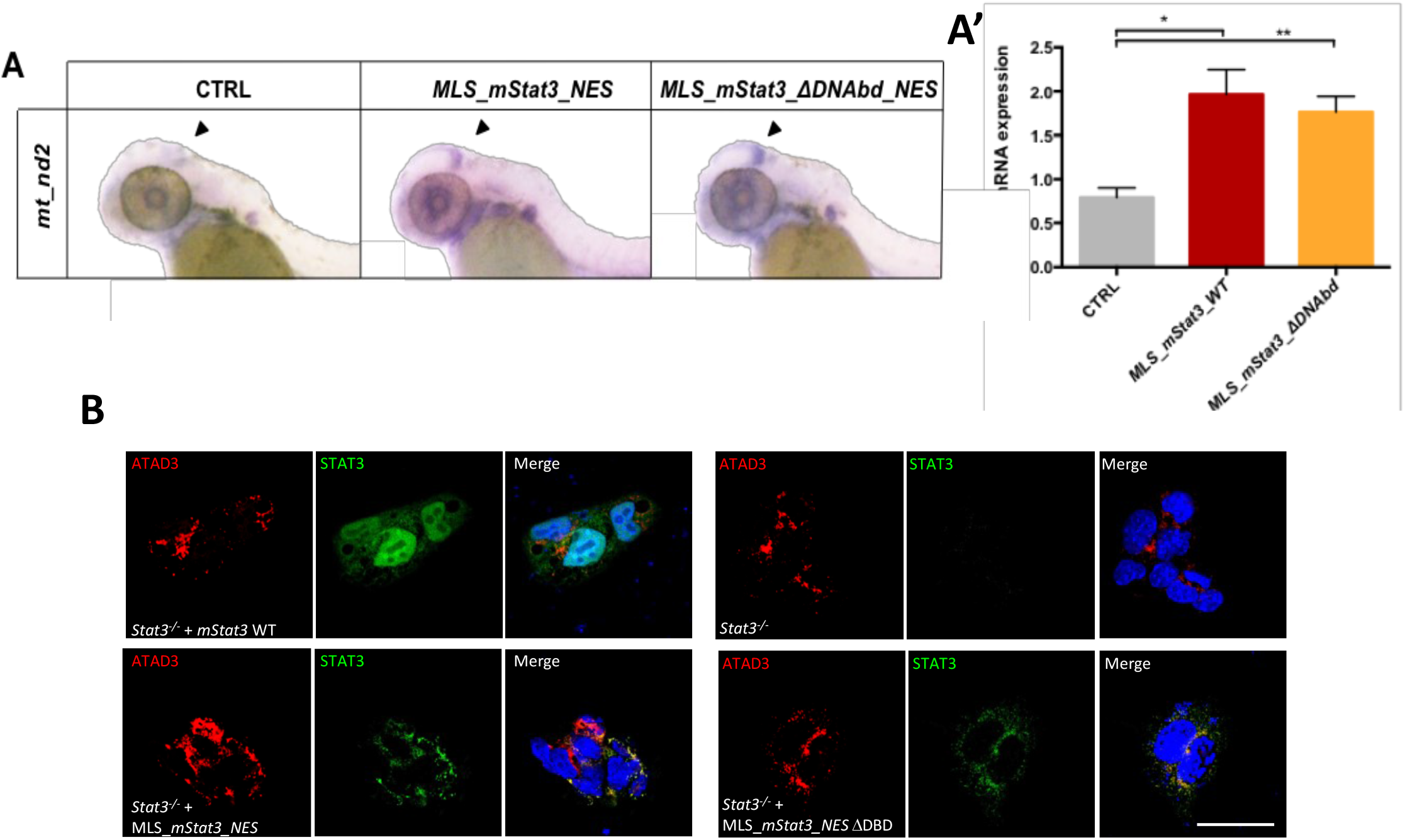
Mutation of Stat3 DBD of does not affect its mitochondrial activities. **A-A’:** qRT-PCR showing *mt_nd2* gene expression after injection of *MLS_mStat3_NES* or *MLS_mStat3_ΔDNAbd_NES* mRNA in 48-hpf embryos; *zgapdh* was used as internal control (p-values= 0.0184; 0.0093). **B:** immunofluorescence on ESCs transiently transfected with *MLS_mStat3_NES* or *MLS_mStat3_ΔDNAbd_NES* and stained with anti-STAT3 (green), anti-ATAD3 (red) Ab and DAPI (blue). Scale bar: 200 μm. Statistical analysis in C-F was performed by unpaired t-test on 3 independent biological samples (where n not specified). ns: not significant; *p<0.05; **p<0.01; error bars=SEM.

The nuclear activity of the STAT3 activation domain is known to be controlled by JAK1/2/3-mediated phosphorylation on Y705 residue, which also ensures STAT3 monomers stability in the cytoplasm (Becker *et al.*, 1998). On the other hand, phosphorylation on STAT3 S727 by the MAPK pathway (Ras-Raf-MEK-ERK pathway) is known, from *in vitro* studies, to enhance the Electron Transport Chain (ETC) (Wegrzyn *et al.*, 2009) as well as to promote cell proliferation and optimal pluripotency (Huang *et al.*, 2014). To verify *in vivo* the post-translational requirements, and to test whether also mitoSTAT3 activity requires Y705 phosphorylation, we decided to inject 1-cell stage zebrafish embryos with mRNAs encoding variants of murine *Stat3.* Specifically, we compared the activity of WT STAT3 (without the MLS) with two mutated forms, Y705F and S727A, able to prevent phosphorylation of residues 705 and 727, respectively (Mohr et al., 2013; Huang et al., 2014). Interestingly, when embryos were injected with the WT isoform, quantitative analysis of fluorescent *in situ* hybridization revealed a significant increase of mitochondrial transcription in the PML of the TeO (Fig. 4 A,B). qRT-PCR analysis on homogenized embryos detected an increase of global *mt_nd2* gene expression (Fig. 4 C,D), which failed to reach statistical significance. Notably, when injecting either Y705F or S727A isoforms of STAT3 no stimulation of mitochondrial transcription in the PML population was detected, either using *in situ* hybridization or qRT-PCR (Fig. 4 A-D; Fig. S4 A). In conclusion, both phosphorylations are needed for STAT3-mediated increase of mtDNA transcription in the PML. On the other hand, when the mutated isoforms are forcedly targeted only to the mitochondrion (by using both the MLS and the NES), the S727A mutation prevented mitoSTAT3-mediated activation of *mt_nd2* gene expression, while the STAT3-Y705F mutated isoform retained its mitochondrial transcriptional activity (Fig. 4 E,F; Fig. S4 B). This implies that Y705 is not directly involved in mitochondrial transcription

**Fig. 4:**
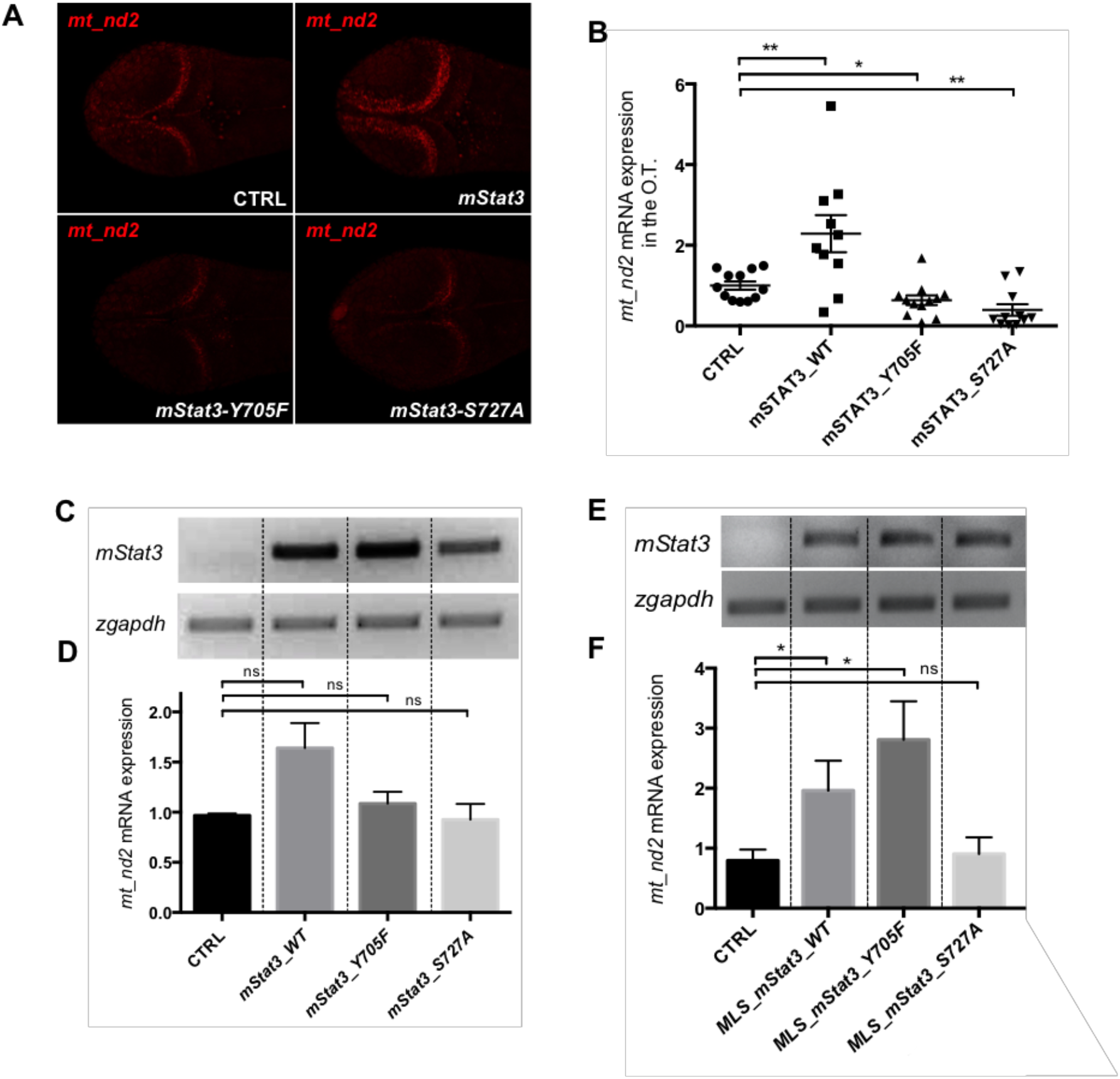
mitoSTAT3 transcriptional activity relies on both S727 and Y705 phosphorylations. **A:** FISH with *mt_nd2* probe in the TeO of 48-hpf embryos injected with mRNA encoding the indicated isoforms of *mStat3.* **B:** Fluorescence quantification of *mt_nd2* mRNA expression in the TeO (n=10) (p-values= 0.0074; 0.0307; 0.0023). **C:** RT-PCR analysis of *mStat3* transcripts detected at 48 hpf/hpi in embryos injected with the indicated form of *mStat3* mRNA; *zgapdh* was used as internal control. **D:** qRT-PCR analysis of *mt_nd2* transcript levels at 48 hpf/hpi normalized to *zgapdh* (p values= 0.0888; 0.1899; 0.8334). **E:** RT-PCR analysis of *MLS_mStat3_NES* transcripts detected at 48 hpf/hpi in embryos injected with indicated form of mitochondria-targeted *mStat3* mRNA; *zgapdh* was used as internal control. **F:** qRT-PCR analysis of *mt_nd2* transcript levels at 48 hpf/hpi normalized to *zgapdh* (p values= 0.0184; 0.0355; 0.5846). Statistical analysis was performed by unpaired t-test on 3 independent biological samples (where n not specified). ns: not significant; *p<0.05; **p<0.01; error bars=SEM.

Given that STAT3 Y705F had no direct effect on mitochondrial transcription (Fig. 4), we hypothesised that the tyrosine 705 could regulate STAT3 localization. To further elucidate the localization of mutated STAT3, we performed immunofluorescence analysis on mouse *Stat3^-/-^* ESCs transiently transfected. While transient transfection of STAT3-Y705F resulted in its nuclear and sporadic mitochondrial localization, transfected mitoSTAT3-Y705F localised exclusively to mitochondria (Fig. 5 A). Surprisingly, upon isolation of mitochondrial fractions followed by western blot analysis we could detect STAT3-Y705F, as well as WT STAT3, in mitochondria (Fig. 5 B-C). Nonetheless, transmission electron microscopy (TEM) analysis after DAB (3,3′-Diaminobenzidine) immunohistochemistry revealed that, while WT STAT3 and MLS_STAT3_NES localize inside mitochondria, staining cristae of the inner mitochondrial membrane (Fig. 5 D), STAT3 Y705F forms clots along the edges of mitochondria and displays a diffuse cytoplasmic signal, fails to migrate through the outer mitochondrial membrane and intermembrane space, thus confirming that Y705 is essential for the correct localization of STAT3 inside the mitochondrion.

**Fig. 5:**
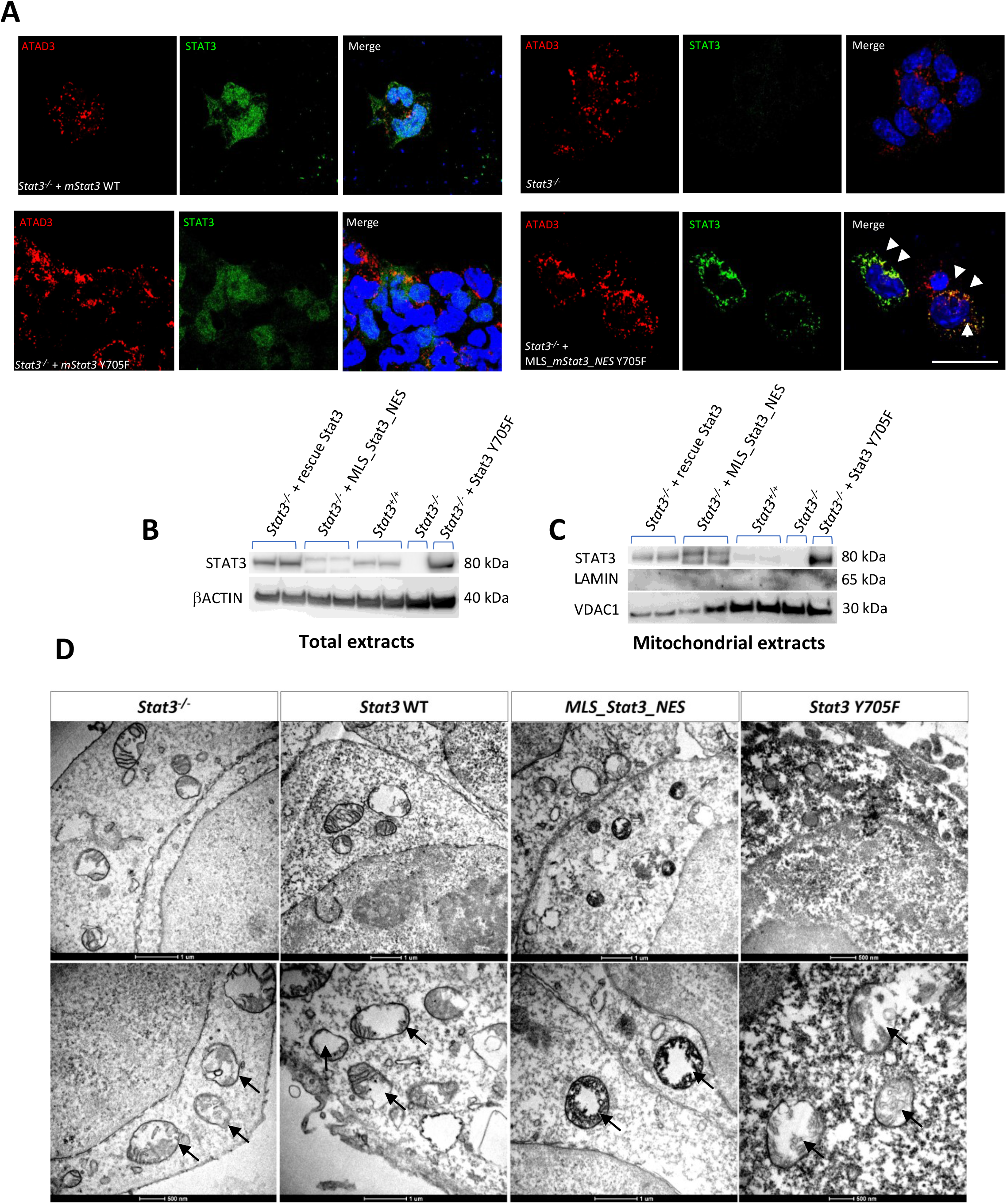
Y705 phosphorylation is needed for the correct localization of STAT3 in the mitochondrion. **A:** immunofluorescence with anti-STAT3 and anti-ATAD3 Ab on ESCs transient transfected with either *mStat3, mStat3 Y705F* or *MLS_mStat3_NES Y705F.* Arrows indicate the colocalization of ATAD and STAT3. Scale bar: 200 μm. **B:** western blot of total STAT3 in ESCs extracts, ß-actin was used as a loading control. **C:** western blot of mitochondrial STAT3 from ESCs mitochondrial extracts; VDAC1 was used as a mitochondrial loading control, Lamin was used as a control for nuclear contamination. **D:** representative pictures of DAB immunohistochemistry on ESCs acquired with TEM; positive signal is black and negative is white. Arrows indicate mitochondria. Cristae are positive in *Stat3* WT and *MLS_Stat3_NES* transfected cells.

### STAT3 S727 phosphorylation is needed for mitoSTAT3-driven promotion of cell proliferation in the PML

As phosphorylation to STAT3 S727 is needed for mitoSTAT3-driven mtDNA transcription, we tested whether this post-transcriptional modification is also required for the increase of PML proliferation downstream of mitochondrial RNA production in the PML.

Indeed, the proliferation rate in the PML of 48-hpf embryos injected with *MLS_mStat3_NES_S727A* mRNA resulted significantly lower to that of embryos injected with WT *MLS_mStat3_NES* mRNA (Fig. 6 A), indicating that S727 phosphorylation is crucial for proliferation in the PML. Moreover, we performed immunofluorescence and western blot analysis on Stat3-/- mouse ESCs transiently expressing either WT STAT3, STAT3 S727A, or MLS_STAT3_NES, showing that mitochondrial localization of STAT3 is not affected by the S727A mutation (Fig. 6 B,C).

**Fig. 6:**
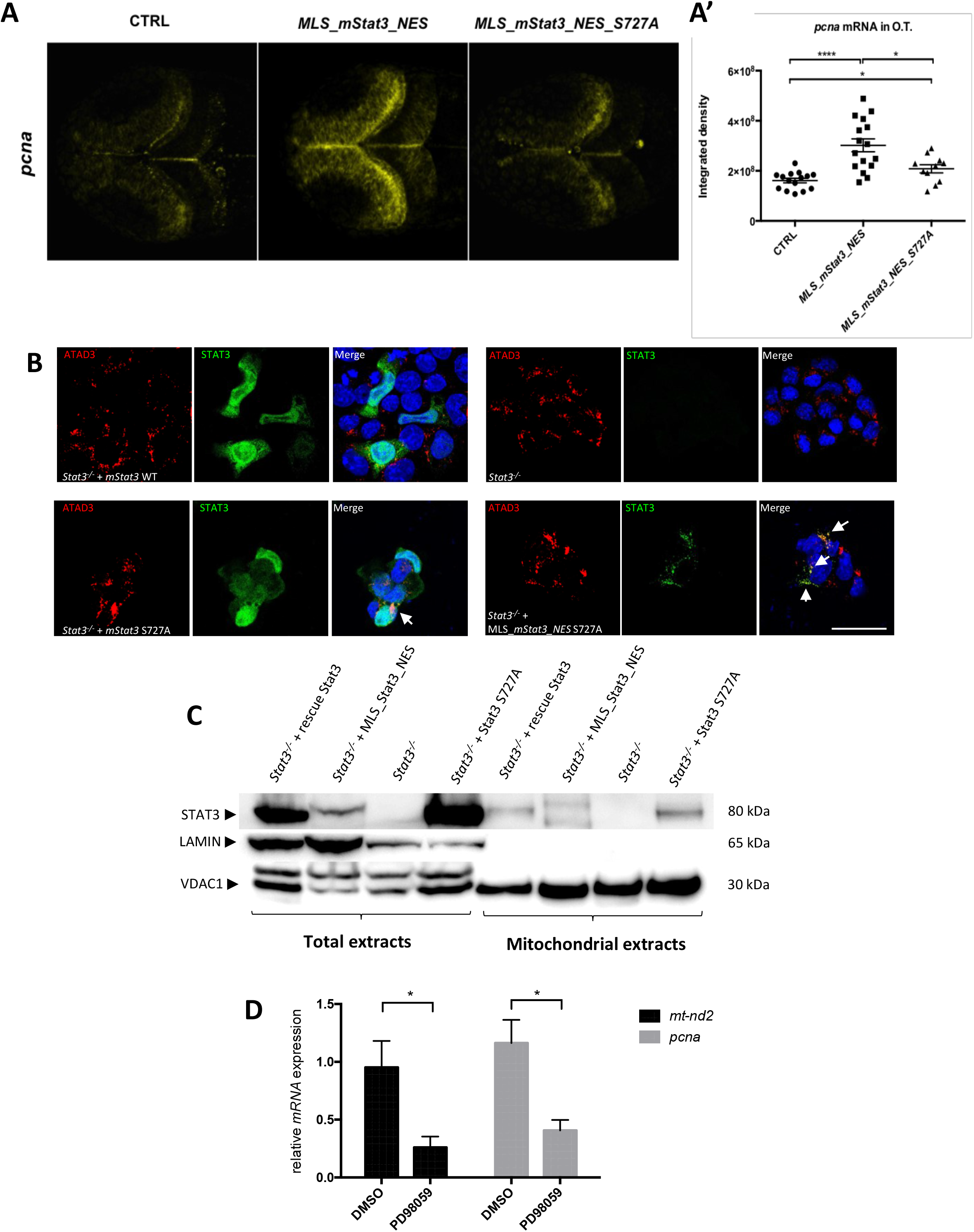
mitoSTAT3-dependent activation of cell proliferation in the TeO depends on S727 phosphorylation. **A-A’:** Representative pictures of WISH performed with an anti-*pcna* probe on 48-hpf uninjected larvae, larvae injected with *MLS_mStat3_NES* and *MLS_mStat3_NES S727A* mRNAs (**A**). Fluorescence quantification of *pcna* mRNA expression in the TeO (n=12) (p-values= <0.0001; 0.0108; 0.0122) (**A’**). **B:** immunofluorescence with anti-STAT3 and anti-ATAD3 Ab on ESCs *Stat3* -/- transiently transfected with the constructs encoding: *mStat3, MLS_mStat3_NES, MLS_mStat3_NES S727A* or mStat3. Arrows indicate the colocalization of ATAD and STAT3. Scale bar: 200 μm. **C:** western blot of mitochondrial STAT3 from ESCs mitochondrial extracts; VDAC1 was used as a mitochondrial loading control, Lamin was used as a nuclear loading control. **D:** qRT-PCR analysis of *mt_nd2* and *pcna* on 48-hpf larvae treated with either PD98059 12.5 μM or DMSO. Statistical analysis was performed by unpaired t-test on 3 independent biological samples (where n not specified). *p<0.05; ****p<0.0001; error bars=SEM.

On the other hand, it is known that in mouse 3T3 fibroblasts the MEK-ERK pathway is responsible for phosphorylation of STAT3 S727 (Gough *et al.*, 2013). In order to evaluate *in vivo* the involvement of MAPK pathway in phosphorylation of S727, and its downstream effects for mitoSTAT3-driven cell proliferation, we decided to use the MEK kinases inhibitor PD98059 (Alessi *et al.*, 1995), commonly used *in vitro* to prevent S727 phosphorylation (Tian and Al., 2004; Wang *et al.*, 2019). First, we tested this compound in 3T3 mouse fibroblasts checking the levels of STAT3 pS727 by western blot. Once confirmed that PD98059 downregulates the phosphorylation of S727 in 3T3 cells (Fig. S5 A), we administered this compound to zebrafish embryos. Interestingly, WT larvae treated from 24-48 hpf with PD98059 displayed a significant reduction of *mt_nd2* and *pcna* transcript levels, suggesting that MEK inhibition reduced both mitochondrial gene expression and cell proliferation (Fig. 6 D).

In conclusion, S727 phosphorylation connects mitochondrial transcription with cell proliferation and inhibitors of the MEK-ERK pathway affecting S727 phosphorylation, such as PD98059, abrogate both processes.

### Jak2 kinase maintains normal mtDNA transcription and proliferation in the PML and the intestine

After demonstrating that a) mitoSTAT3-driven mitochondrial transcription relies on both Y705 and S727 post-transcriptional phosphorylation, and b) that the consequential proliferation effect downstream of mitochondrial transcription requires functional MEK kinases, we decided to test the dependence of both mitochondrial transcription and cell proliferation on Jak2 Tyrosine-kinase activity, which promotes Y705 phosphorylation. WT embryos were therefore treated from 24 to 72 hpf with AG490, a specific inhibitor of Jak2 widely used as a JAK/STAT3 inhibitor (Park *et al.*, 2014; Garbuz *et al.*, 2014), and the expression of *mt_nd2* was assessed by qRT-PCR. When observed at 72 hpf, AG490-treated larvae displayed a significant reduction of *mt_nd2* expression in the PML, the inner retina and the primordium of the intestine (Fig. 7 A (arrowheads)), while no significant decrease was present at 48 hpf (Fig. 7 B; Fig. S6 A). In addition, proliferation activity was found to be significantly reduced in the PML of 72-hpf AG490-treated larvae as assayed by *in situ* hybridization using anti-*pcna* probe (Fig. 7 C, D).

**Fig. 7:**
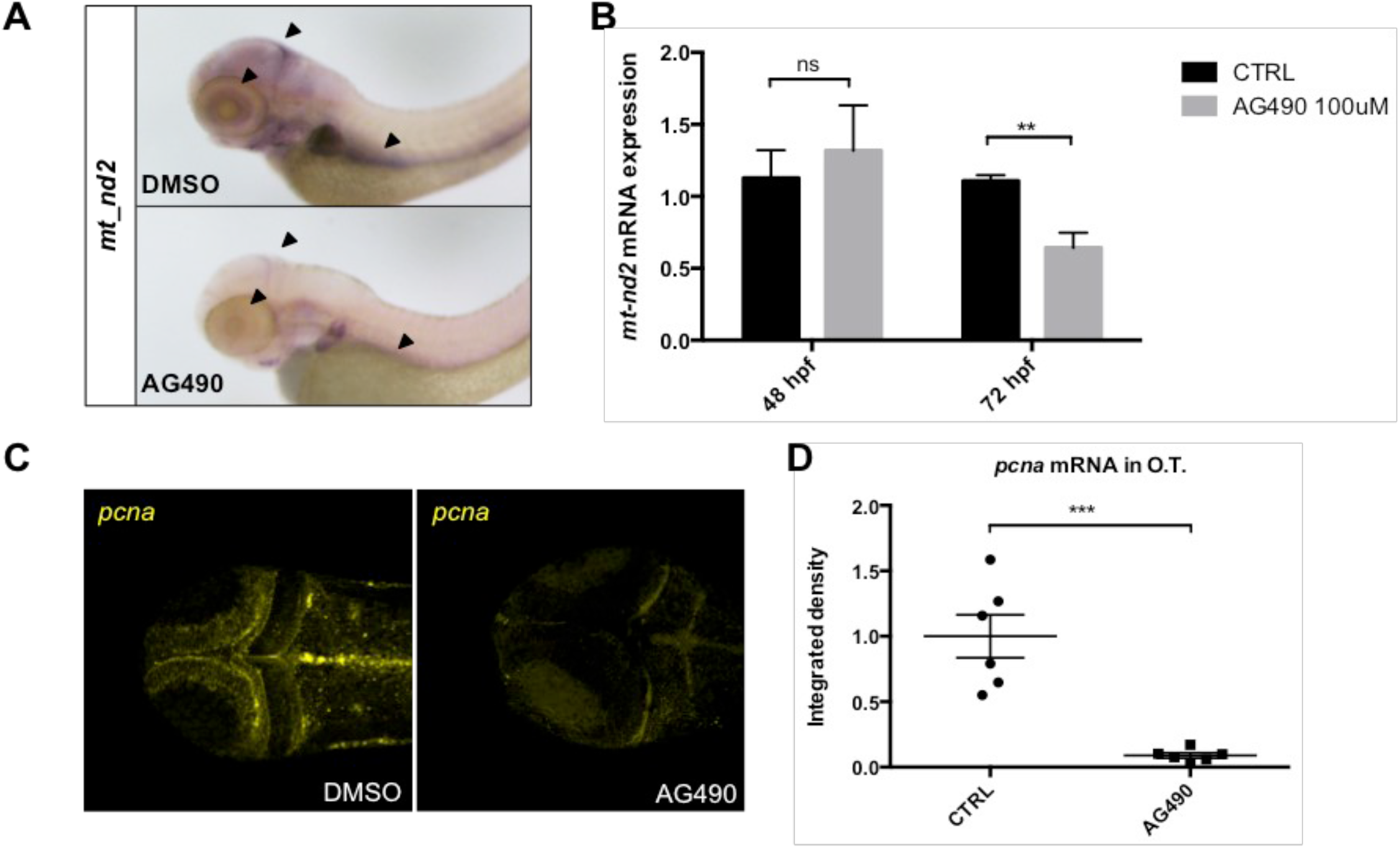
JAK inhibition impairs normal mitochondrial transcription and cell proliferation in the TeO of 72-hpf embryos. **A:** WISH with anti-*mt_nd2* mRNA probe on 72-hpf embryos treated with 100 μM AG490 from 24-72 hpf and DMSO treated controls. **B:** relative *mt_nd2* transcript expression assayed by qRT-PCR in 48- and 72-hpf embryos treated with 100 μM AG490 and DMSO treated controls starting from 24 hpf; *zgapdh* was used as internal control (p-values= 0.6261; 0.0060). **C:** FISH with anti-*pcna* probe in the TeO of 72-hpf embryos treated with 100 μM AG490 from 24 to 72 hpf and DMSO treated controls. **D:** Fluorescence quantification of *pcna* mRNA expression in the TeO (n=6) (p value=0.0003). Statistical analysis was performed by unpaired t-test on 3 independent biological samples (where n not specified). ns: not significant; **p<0,01; ***p<0.001; error bars=SEM.

We also investigated the effect of Jak2 inhibitor in the intestine, a highly proliferating tissue of zebrafish larvae, where *mt_nd2* gene is strongly expressed between 3 and 6 days post fertilization (dpf) (Fig. 8 A). The activity of Stat3 in the intestine of zebrafish is consistent with the facts that a) the proliferative and survival effects of IL-6 in murine IECs (intestinal epithelial cells) is largely mediated by STAT3 (Grivennikov *et al.*, 2009), b) that STAT3 is needed for small-intestine crypt stem cell survival, as revealed by conditional mutant mice (Matthews *et al.*, 2011), and c) that Stat3-positive cells in zebrafish intestine represent a population of intestinal Wnt-responsive stem cells (Peron *et al.*, 2020). Administration of 60 μM AG490 between 3 and 6 dpf was able to significantly reduce mitochondrial transcription in the intestine of treated larvae with respect to DMSO treated controls (Fig. 8 A, B). Moreover, the treatment of larvae with AG490 caused a significant decrease in the number of intestinal proliferating cells (revealed by immunohistochemistry with anti-pH3 antibody) (Fig. 8 C, D) and resulted in flattening of the intestinal mucosa (Fig. 8 E, F). Taken together, these experiments demonstrate *in vivo* that phosphorylation of the Stat3 Y705 residue is required in zebrafish for normal mitochondrial transcription and downstream proliferation in the developing TeO and intestine.

**Fig. 8:**
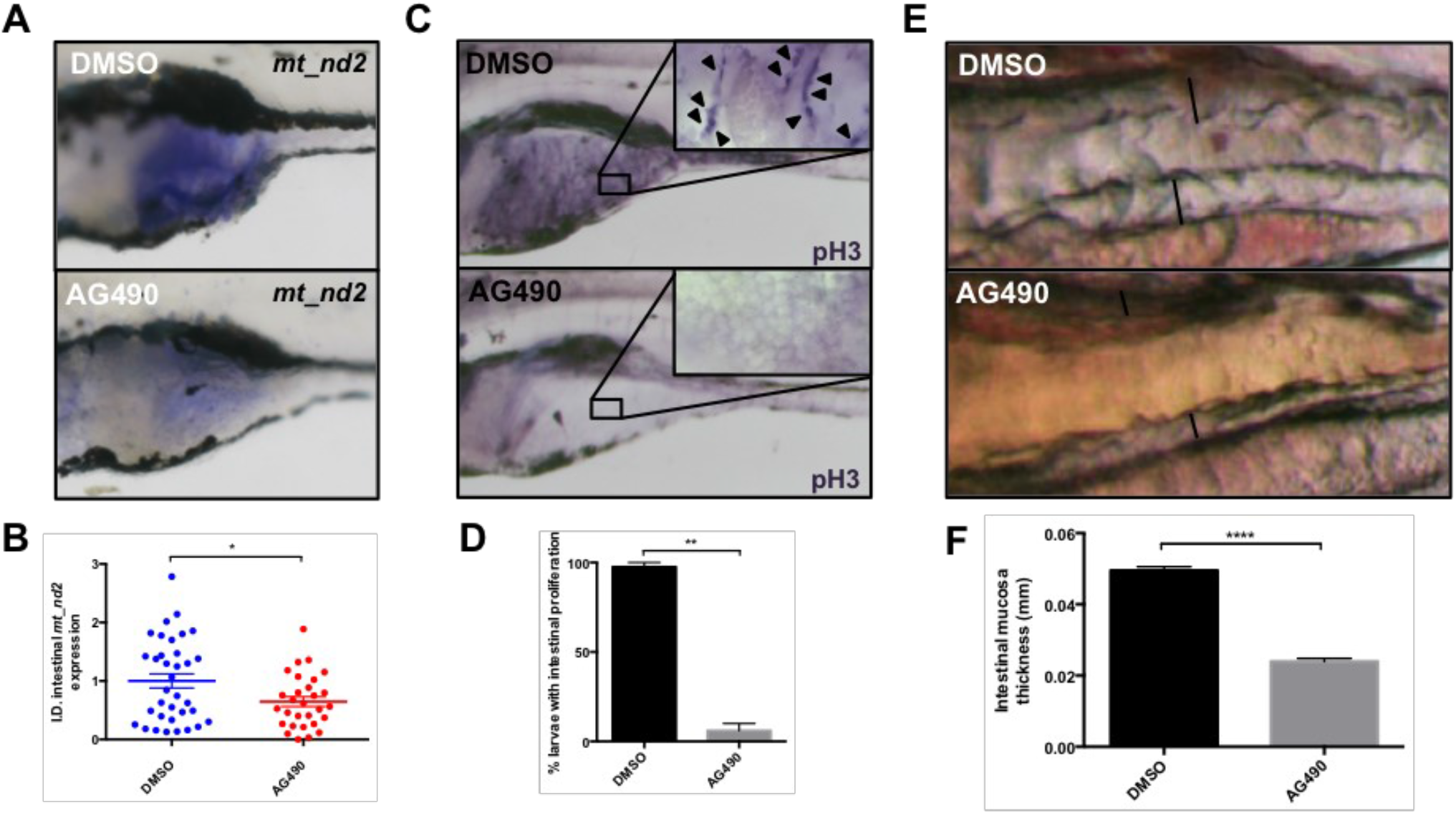
JAK inhibition impairs normal mitochondrial transcription and cell proliferation in the intestine of 6-dpf larvae. **A:** WISH with *anti-mt_nd2* mRNA probe on 6-dpf larvae treated with 60 μM AG490 from 24-72 hpf and DMSO treated controls; zoom on the intestine. **B:** Quantification of *mt_nd2* mRNA expression in the intestine (n=30) (p-value= 0.0240). C: phospho-Histone-H3 (pH3) immunostaining of 6-dpf AG490 treated larvae and DMSO treated controls; zoom on the intestine; (pH3 positive cells=arrowheads). **D:** Quantification of the number of AG490 and DMSO treated larvae displaying intestinal proliferation (n=15) (p-value= 0.0026). **E:** AG490 treated larvae showing loss of folding in intestinal mucosa. **F:** Graph showing the dimension of mucosal thickness in both DMSO and AG490 6-dpf treated larvae (n=18) (p-value= 0,0001). Statistical analysis was performed by unpaired t test on indicated number of samples; *p<0.05; **p<0.01; ***p<0.001; error bars=SEM.

### The zebrafish *stat3^-/-^* null mutant displays impairment of mitochondrial transcription and cell proliferation in CNS and intestine

To confirm data obtained by endogenous Stat3 chemical treatment with either MEK and Jak2 inhibitors, we used the zebrafish *stat3^ia23^* mutant (from now on called *stat3^-/-^* (Peron *et al.*, 2020), which is predicted to encode a premature stop codon at amino acid 456, thus lacking all functional domains including the dimerization domain and the transactivation domain, harbouring Y705 and S727 phosphorylation sites, respectively. As reported in Peron *et al.* (2020), these mutants die within one month of age and they can be obtained only after breeding between adult *stat3^+/-^* zebrafish. We decided to test if the genetic ablation of zebrafish *stat3* determines a reduction of *mt_nd2* in 48-hpf larvae. As reported in Fig. S6 B, no significant differences were detected by *in situ* hybridization against *mt_nd2* in *stat3^+/+^, stat3^+/-^*, and *stat3^-/-^* 48-hpf sibling larvae. This result is probably due to genetic compensation. To overcome this issue and to determine whether genetic ablation of *stat3* alters mitochondrial transcription and cell proliferation in 48-hpf larvae, we decided to analyse *stat3* “CRISPants” generated after injection of Cas9 protein and sgRNAs which target the antisense strand of exons 14, 22 and 23 of *stat3* gene: this approach is employed to target a gene and silence its RNA transcription, according to the principles of classic CRISPR/Cas9 (Strutt et al., 2018). We evaluated the efficiency of *stat3* targeting by injection of the *Tg(7xCRP-Hu.tk:EGFP)* reporter line characterized in Peron *et al.* (2020). Results show that Stat3-dependent fluorescence of reporters is significantly dampened in CRISPants when compared to control larvae (Fig. 9 A-A’), while PCR analysis of the target confirmed that *stat3* has been successfully mutagenised in exons 14, 22, and 23 (Fig. 9 A’’). Notably, qRT-PCR analysis of *stat3* and *socs3a* in CRISPants displayed a significant downregulation of both transcripts (Fig. 9 B,D). Hence, we decided to use CRISPants for the analysis of *mt_nd2* and *pcna* expression levels. As reported in Fig. 9 E-F, both transcripts are significantly downregulated in *stat3* CRISPants, confirming again that Stat3 is involved in mitochondrial transcription and cell proliferation. Interestingly, injection of *MLS_mStat3_NES* mRNA in CRISPants, while rescuing completely *mt_nd2* and *pcna* transcript levels, does not restore the nuclear activities of Stat3 (Fig. 9 C,D,E,F).

**Fig. 9:**
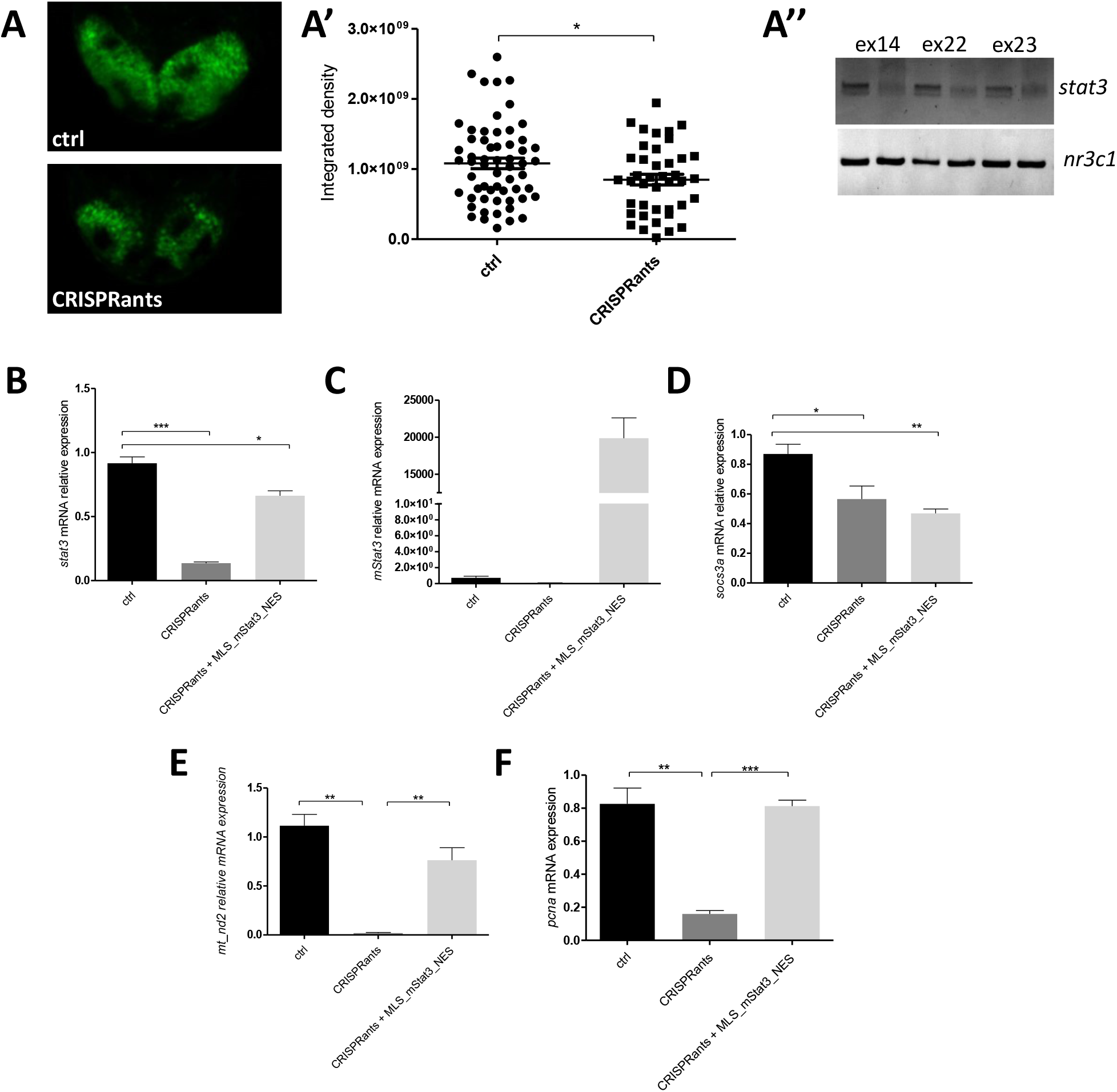
*stat3* CRISPRants show reduced mitochondrial transcription that is rescued by mitochondrial *Stat3*. **A-A’’:** Representative pictures of 48-hpf *Tg(7xStat3:EGFP)* transgenic zebrafish larvae. Fluorescent quantification of TeO of control and CRISPRant zebrafish larvae. PCR amplification of the *stat3* gene on DNA extracts from control and CRISPRant larvae *(nr3c1* gene is used as an internal control). **B:** qRT-PCR against *stat3* in 48-hpf control, CRISPRants and CRISPRants + *MLS_mStat3_NES* mRNA zebrafish larvae. **C:** qRT-PCR against *Stat3* in 48-hpf control, CRISPRants and CRISPRants + *MLS_mStat3_NES* mRNA zebrafish larvae. **D:** qRT-PCR against *socs3a* in 48-hpf control, CRISPRants and *CRISPRants+MLS_mStat3_NES* mRNA zebrafish larvae. **E:** qRT-PCR against *mt_nd2* in 48-hpf control, CRISPRants and *CRISPRants+MLS_mStat3_NES* mRNA zebrafish larvae. **F:** qRT-PCR against *pcna* in 48-hpf control, CRISPRants and *CRISPRants+MLS_mStat3_NES* mRNA zebrafish larvae. Statistical analysis was performed by unpaired t-test on 3 independent biological samples (where n not specified). *p<0.05; **p<0.01; ***p<0.001; error bars=SEM.

In agreement with our previous results, when analysed at 6 dpf *stat3* knock-out displays a significant and clear reduction of both *mt_nd2* and *pcna* transcripts, endorsing the link between Stat3 mitochondrial functions and its role in the regulation of cell proliferation. Notably, about 70% of *stat3^-/-^* larvae display severe defects in the development of intestinal epithelium (Peron *et al.*, 2020): as revealed by pHH3 immunostaining, intestinal mitoses are almost absent (Fig. 10 B-B’) and the intestine fails to fold (Fig. 10 C-C’). These phenotypic alterations are almost identical to those induced by AG490 treatment (Fig. 8 B,C). At 6 dpf *stat3^-/-^* larvae also show impaired CNS cell proliferation in the Telencephalon (Tel), the Diencephalon (Di) and the TeO, where Pcna is found to be reduced down to 15% with respect to *stat3^+/+^* siblings, supporting, once again, the requirement of Stat3 to maintain normal proliferation in the brain (Fig. 10 D, E).

**Fig. 10:**
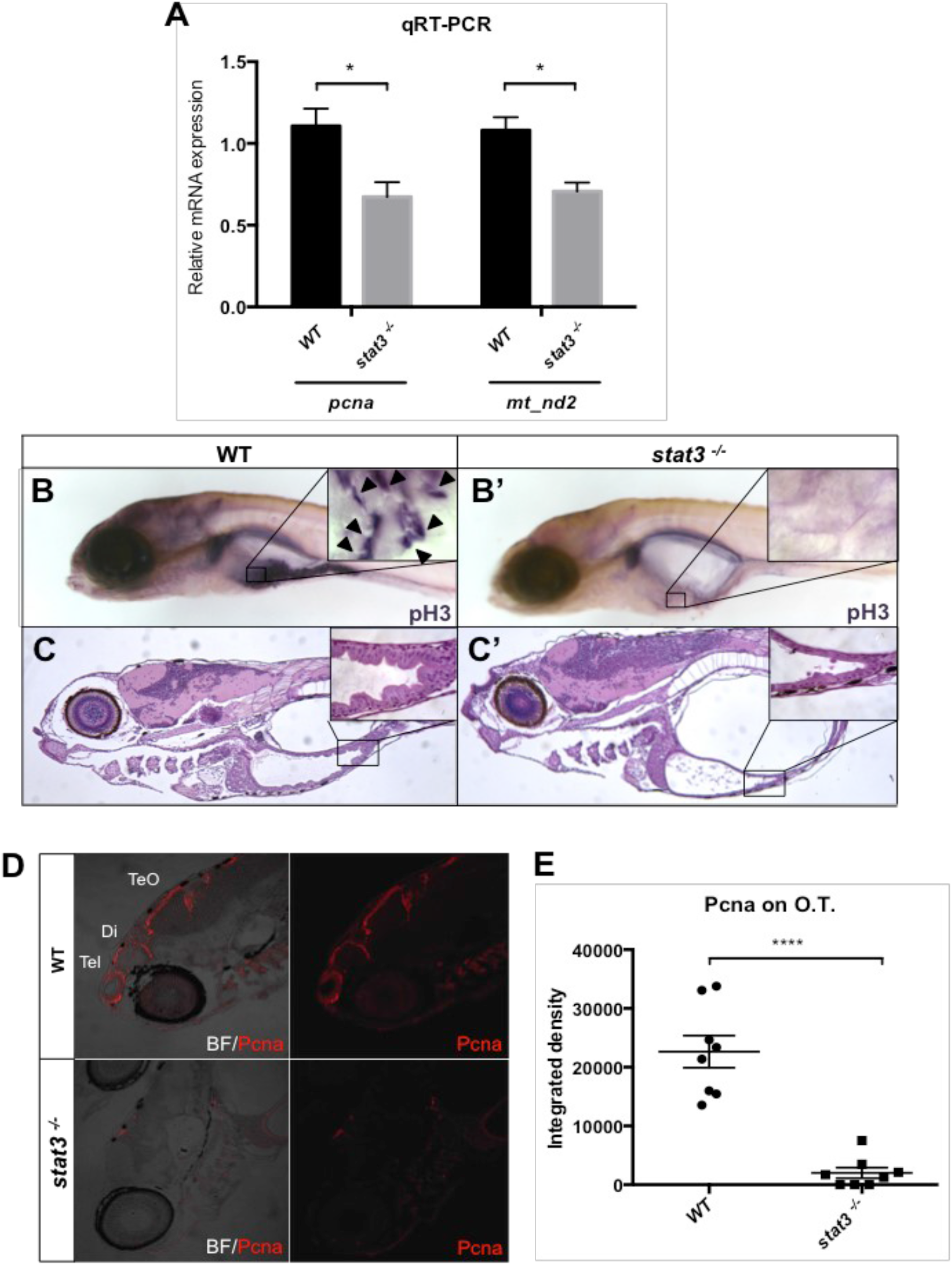
*stat3* KO impairs normal mitochondrial transcription and cell proliferation in the intestine and brain of 6-dpf zebrafish larvae. **A:** Relative mRNA expression of *mt_nd2* and *pcna* transcripts assayed by qRT-PCR in homogenized *stat3^-/-^* and WT siblings at 6 dpf; *zgapdh* was used as internal control (p values= 0.0358; 0.0182). **B-B’:** phospho-Histone-H3 (pH3) immunostaining of *stat3^-/-^* and WT siblings at 6 dpf; zoom on the intestine. (pH3 positive cells=arrowheads). **C-C’:** EE staining on WT and *stat3^-/-^* mutant sections at 6 dpf shows the complete loss of folding in the mutant intestinal epithelium. **D:** IF with anti-PCNA Ab on 6-dpf *stat3^-/-^* mutants showing decrease of fluorescence in the CNS (Tel= telencephalon; Di: diencephalon; TeO: tectum opticum). **E:** Fluorescence quantification of PCNA protein on lateral sections of 6-dpf *stat3^-/-^* mutants and WT siblings; zoom on the head (n=8) (p-value<0.0001). Statistical analysis was performed by unpaired t-test on 3 independent biological samples (where n not specified). *p<0.05; ****p<0.0001; error bars=SEM.

Interestingly, no overt structural alteration is present in intestinal or brain mitochondria of 6-dpf *stat3^-/-^* larvae analysed by TEM (Fig. S7 A). Moreover, in order to evaluate the amount of mitochondria, we crossed *stat3* mutants with the *Tg(CoxVIII-mls:EGFP)* transgenic line that expresses a mitochondria-localized form of enhanced GFP. No clear change in the total mitochondria volume was present in the intestine of *stat3^-/-^* larvae with respect to *stat3^+/+^* sibling larvae (Fig. S7 B,C). Together with previous results, this highlights that mitoSTAT3 is only acting as regulator of mitochondrial transcription, without impacting on mitochondria biogenesis or homeostasis.

## DISCUSSION

Using the zebrafish model and taking advantage of a STAT3 harbouring both a mitochondrial localization sequence and a nuclear export signal, we explored how mitoSTAT3 may act inside the mitochondrion. Interestingly, since mitochondrial mRNAs a) are reduced in *stat3^ia23/ia23^* zebrafish null mutants and CRISPants b) are decreased in embryos treated with the Jak2 kinase inhibitor AG490, and c) the effect of *MLS_Stat3_NES* in promoting mitochondrial gene expression is abolished by Balapiravir (a mtRNA RNA polymerase inhibitor), our data strongly support, *in vivo*, a direct link between mitoSTAT3 activity and mitochondrial transcription. This is consistent with the mitochondrial transcriptional role of mitoSTAT3 found *in vitro* in murine ESCs, previously reported (Carbognin *et al.*, 2016). On the other hand, quite surprisingly for a transcription factor, *MLS_Stat3_NES* mutated in its DNA-binding domain is still able to increase mitochondrial transcription. This result suggests that STAT3, differently from what hypothesized in Macias *et al.* (2014), does not regulate mtDNA transcription by binding STAT3 responsive elements located in the mtDNA, consistently with the differences between the eukaryotic and the prokaryotic transcriptional machineries operating in the nucleus and mitochondria, respectively. We previously reported that Stat3 binds mtDNA in ESCs (Carbognin *et al.* 2016), therefore we hypothesise that such binding is mediated by additional proteins whose identification will be the aim of future studies.

One of the most fascinating aspects of mitochondria evolution is their progressive incorporation in the machinery of cell regulatory activities such as cell proliferation and apoptosis (Antico Arciuch *et al.*, 2012). By showing that mitoSTAT3-driven mitochondrial transcription controls cell proliferation, at least in intestinal and tectal undifferentiated progenitor cells, our data partially answer the open questions about the mechanisms that synchronize mitochondrial and nuclear activities during cell proliferation.

Canonical STAT3 activation depends on different modifications, such as the phosphorylation at tyrosine 705 (Y705), that induces dimerization and translocation to the nucleus, and at serine 727 (S727), whose function has been reported to have unclear effects on STAT3 nuclear transcriptional activity (Decker *et al.*, 2000; Huang *et al.*, 2014). On the other hand, the post-translational modifications required for mitoSTAT3 import and activity in mitochondria have not been clearly dissected so far, although phosphorylation at S727 has been found to both activate OXPHOS complexes I and II, and suppress ROS production and cytochrome c release following ischemic injury (Meier & Larner, 2014). More recently, STAT3 phosphorylation at S727 was also found to be required for STAT3-mediated regulation of ER Ca^2+^ fluxes and apoptosis through the regulation of the mitochondrial Ca^2+^ uptake (Avalle *et al.*, 2019). We provide here *in vivo* evidence that phosphorylation of STAT3 Y705, being required for precise mitochondrial import of STAT3, is needed for STAT3-mediated mitochondrial gene expression, a result also confirmed in mouse ESCs. On the other hand, in accordance with the results obtained by Wegrzyn *et al.* (2009), we show that mitochondrial STAT3 transcriptional activity *in vivo* is totally dependent on phosphorylation of the ERK target S727. Experiments in mouse ESCs allowed us to clearly demonstrate that phosphorylated S727 is not required for mitochondrial localization of STAT3 but, rather, for its activity once imported in the organelle. *In vivo* experiments performed in zebrafish larvae show that both mitoSTAT3-mediated mtDNA transcription and cell proliferation are repressed by targeting S727 with a MEK inhibitor.

Hence, by dissecting the roles of Y705 and S727 phosphorylation in the mitochondrial specific activity of STAT3 both in zebrafish larvae and mouse ESCs, our results add further insight into the specificity of mitochondrial STAT3 in the regulation of cellular processes, previously thought to be dependent exclusively on canonical (nuclear) STAT3. Together with the fact that mitochondrial STAT3 has been identified as a contributor to RAS-dependent cellular transformation (Gough *et al.*, 2009), we support the idea of ERK-mitoSTAT3-mediated mitochondrial transcription might be a key process in cancer development, especially in the intestine, where we demonstrate here and in Peron et al., (2020) that cell proliferation is STAT3-dependent. Considering that, to date, the vast majority of STAT3-targeted cancer therapeutic approaches focus only on its canonical functions, our findings imply mitochondrial STAT3-specific transcriptional activity as a significant molecular mechanism to be targeted.

## MATERIALS AND METHODS

### Animal husbandry and lines

Animals were staged and fed as described by Kimmel *et al.* (1995) and maintained in large scale aquaria systems.

Embryos were obtained by natural mating, raised at 28 °C in Petri dishes containing fish water (50X: 25 g Instant Ocean, 39.25 g CaSO_4_ and 5 g NaHCO_3_ for 1 L) and kept in a 12:12 light-dark (LD) cycle. All experimental procedures complied with European Legislation for the Protection of Animals used for Scientific Purposes (Directive 2010/63/EU).

*stat3^ia23^* mutants and *Tg(7xStat3:EGFP)* transgenic zebrafish are described in Peron *et al.* (2020). The *Tg(CoxVIII-mls:EGFP)* transgenic zebrafish line is described in Martorano *et al.* (2019).

### Drug treatments

The following chemical compounds were used: AG490 (T3434, Sigma Aldrich); PD98059 (PHZ1164, Thermo Fisher Scientific); Balapiravir (HY-10443, DBA). Before drug administration, a hole was made in the chorion of 8 hpf embryos, while 24 hpf embryos were dechorionated. All drugs were dissolved in DMSO and stored in small aliquots at −20°C. 100 μM AG490 treatment was performed from 24 to 48 hpf or from 24 to 72 hpf. 60 μM AG490 was administered in 3-6 dpf treatments. 12.5 μM PD98059 treatment was administered from 24 to 48 hpf. 50 μM Balapiravir solution was administered from 8 to 48 hpf. After treatments, embryos were either anesthetized and fixed in 4% paraformaldehyde (PFA) (158127, Sigma) in PBS for ISH, FISH and IHC or in TRI Reagent^®^ (T9424, Sigma) for qRT-PCR analysis.

### CRISPRants generation

sgRNA against exon 14 of *stat3* gene was produced as described in Peron *et al.* (2020). sgRNAs against exon 22 and 23 of *stat3* gene have been designed with CHOPCHOP software https://chopchop.rc.fas.harvard.edu, and provided by Synthego.

Cas9 protein (M0646, NEB) and sgRNAs against antisense of exons 14, 22 and 23 of *stat3* genes were injected in 1-cell stage eggs. Subsequently, 48-hpf injected larvae were collected for DNA and RNA extraction and for imaging. Sequences of sgRNAs and of primer used for genotyping are listed in Table 1.

**Table 1:**
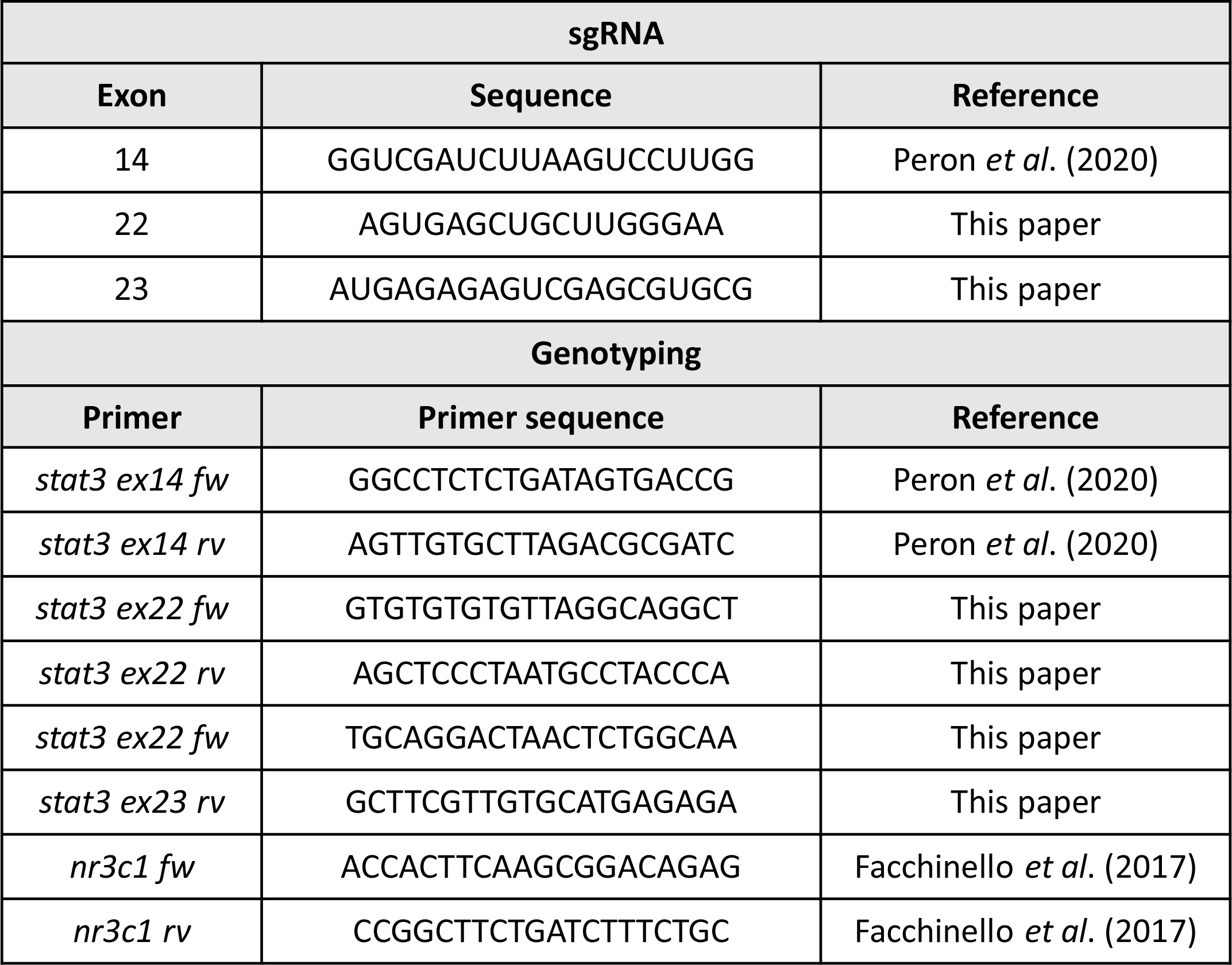
List of sgRNAs and primers used for genotyping (5′-3′ sequences)

### mRNAs synthesis and injection

*mStat3, mStat3_Y705F* and *mStat3_S727A* CDSs were obtained from pCEP4-*Stat3-WT*, *pCEP4-Stat3-Y705F, pCEP4-Stat3-S727A* plasmids (a kind gift of the Poli Lab; Department of Molecular Biotechnology and Health Sciences, Molecular Biotechnology Center, University of Turin) and sub-cloned into a pCS2+ backbone using the In-Fusion^®^ HD Cloning Kit (Clontech). *MLS_mStat3_NES* CDS, containing the murine *Stat3* cDNA flanked by a Mitochondrial Localization Sequence (MLS) and a Nuclear Export Sequence (NES), was subcloned into a pCS2+ plasmid from a 70_pPB-CAG+MLS+*Stat3*+NES-pA-pgk-hph-2-2 plasmid by digestion with *XbaI* and *BamHI.* Mutated forms of *MLS_mStat3_NES* mRNA were obtained from *pCS2+MLS_mStat3_NES* by site-directed mutagenesis using the Q5^®^ Site-Directed Mutagenesis Kit (NEB); primers are indicated in Table 2.

**Table 2:**
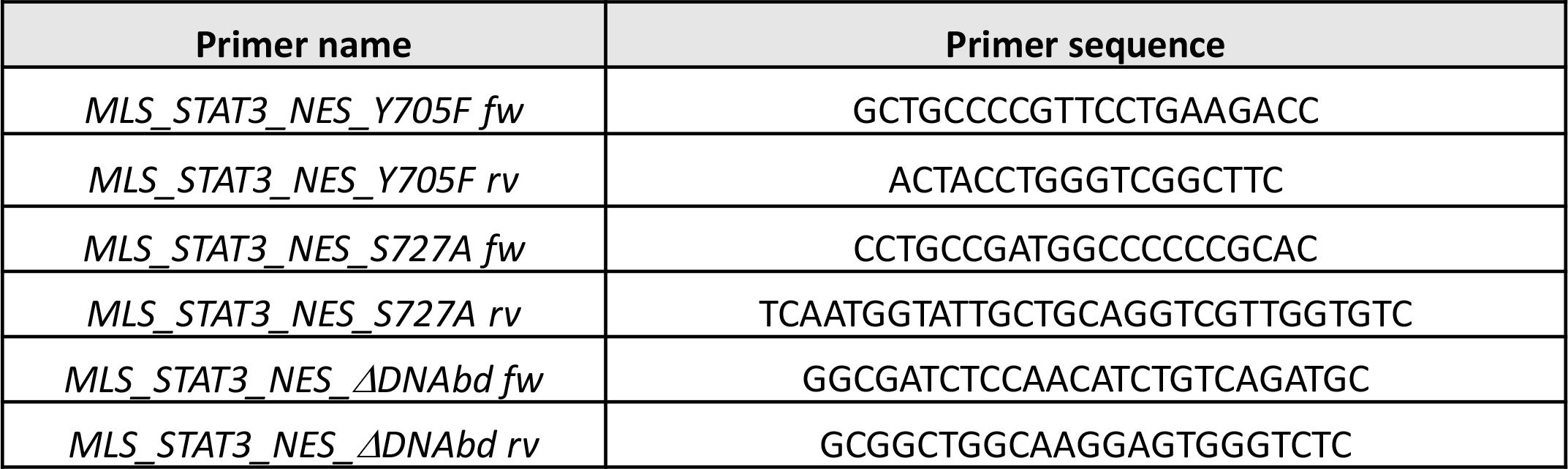
List of cloning-related primers (5′-3′ sequences)

mRNAs were *in vitro* transcribed using the mMESSAGE mMACHINE^®^ SP6 Transcription Kit (Thermo Fisher Scientific) and purified using the RNA Clean and Concentrator kit (Zymo Research). A mix containing mRNA (30 ng/μL for *Stat3-WT, Stat3-Y705F, Stat3-* S727A; 50 ng/μL for *MLS_Stat3_NES)*, Danieau injection Buffer and Phenol Red injection dye, was injected into 1-cell stage embryos.

### mRNA isolation and quantitative real time reverse transcription PCR (qRT-PCR)

For expression analysis, total RNA was extracted from pools of 15 7-dpf larvae or 35 48-hpf embryos with TRIzol reagent (15596018, Thermo Fisher Scientific). mRNA was treated with RQ1 RNase-Free DNase (M6101, Promega) and then used for cDNA synthesis with Superscript III Reverse Transcriptase (18080-044, Invitrogen) according to the manufacturer’s protocol. qPCRs were performed in triplicate with EvaGreen method using a Rotor-gene Q (Qiagen) and the 5x HOT FIREPol ^®^ EvaGreen^®^ qPCR Mix Plus (08-36-00001, Solis BioDyne) following the manufacturer’s protocol. The cycling parameters were: 95 °C for 14 min, followed by 45 cycles at 95 °C for 15 s, 60 °C for 35 s, and 72°C for 25 s. Threshold cycles (Ct) and dissociation curves were generated automatically by RotorGene Q series software. Sequences of specific primers used in this work for qRT-PCR and RT-PCR are listed in Table 3. Primers were designed using the software Primer 3 (http://bioinfo.ut.ee/primer3-0.4.0/input.htm). Sample Ct values were normalized with Ct values from zebrafish *gapdh* and results were obtained following the method described in Livak and Schmittgen (2001).

**Table 3:**
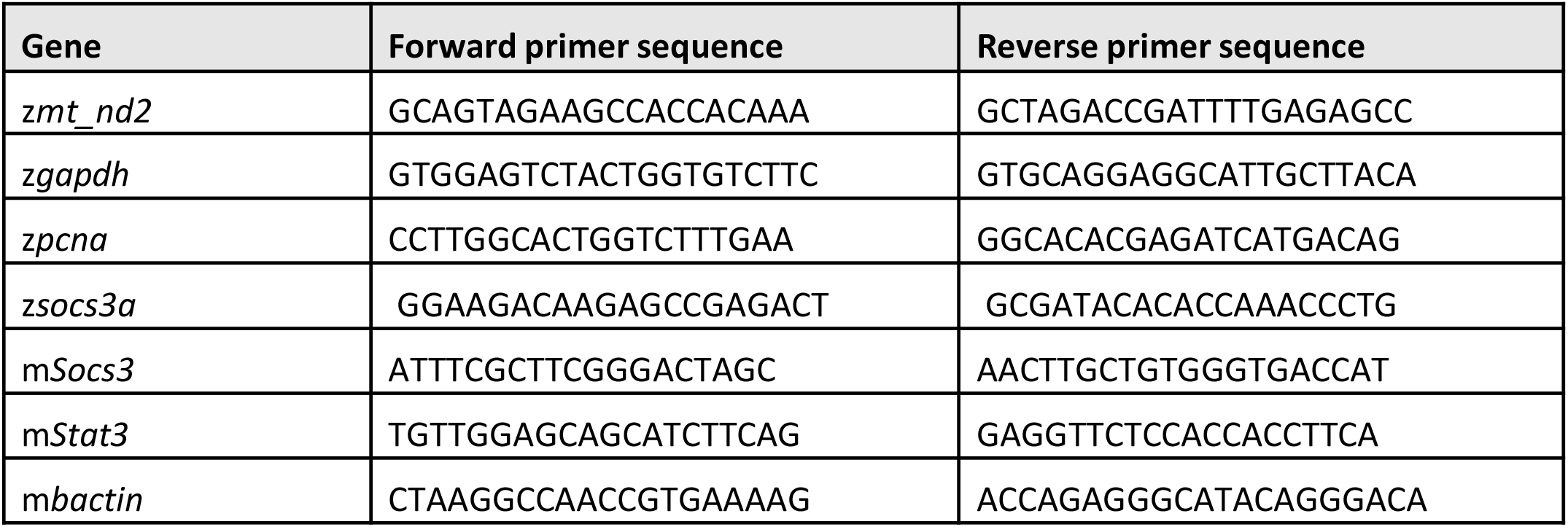
List of qRT-PCR and RT-PCR primers (5′-3′ sequences)

### Immunoblotting and mitochondria isolation

Immunoblotting was performed as previously described in Carbognin *et al.* (2016). The following antibodies were used: anti-STAT3 mouse monoclonal (9139, Cell Signalling) (1:1000), anti-GAPDH mouse monoclonal (MAB374, Millipore) (1:1000), anti-VDAC1 rabbit polyclonal (ab15895, Abcam) (1:1000), anti-Lamin (sc-6217, Santa Cruz) (1:1000), anti-bActin mouse monoclonal (MA1-744, Invitrogen) (1:10000), anti-pSTAT3 S727 rabbit monoclonal (9134, Cell Signalling). Mitochondria from mouse ESCs were isolated using Mitochondria isolation kit (89874, Thermo Scientific).

### 3,3′-Diaminobenzidine staining

Cells were fixed in a 24 wells plate with 4% Paraformaldehyde in PBS (pH 7.4) for 30 minutes at RT (room temperature). After fixation cells were washed 5 times with PBS (5 minutes each), blocked and permeabilized with 5% normal goat serum and 0.1% saponin in PBS for 30 min, and then incubated with primary antibody anti-STAT3 mouse monoclonal (9139, Cell Signalling) ON at 4°C in PBS 5% normal goat serum and 0.05% saponin. After 5 washes with PBS (5 minutes each), cells were incubated with HRP-conjugated Fab fragments of the secondary antibody for 2 hours at RT. After 5 washes, cells were incubated in the DAB solution (0.01 gr DAB in 20 mL TRIS-HCl buffer plus 30% H2O2 solution just before use). Subsequently the samples ware postfixed with 1% osmium tetroxide plus potassium ferrocyanide 1% in 0.1 M sodium cacodylate buffer for 1 hour at 4°C. After three water washes, samples were dehydrated in a graded ethanol series and embedded in an epoxy resin (Sigma-Aldrich). Ultrathin sections (60-70 nm) were obtained with an Ultrotome V (LKB) ultramicrotome, counterstained with uranyl acetate and lead citrate and viewed with a Tecnai G2 (FEI) transmission electron microscope operating at 100 kV. Images were captured with a Veleta (Olympus Soft Imaging System) digital camera.

### Immunofluorescence

ESCs were grown and transfected as described by Carbognin *et al.* (2016). For IF ESCs were fixed for 10 min in cold methanol at −20 °C, washed in TBS, permeabilized for 10 min with TBST + 0.3% Triton X-100 at RT, and blocked for 45 min in TBS + 3% goat serum at RT. The cells were incubated overnight at 4 °C with primary antibodies (anti-STAT3 mouse monoclonal (9139, Cell Signalling) (1:100); anti-ATAD3A rabbit monoclonal (224485, AB-Biotechnologies) (1:100). After washing with TBS, the cells were incubated with secondary antibodies (Alexa, Life Technologies) for 30 min at RT. Cells were mounted with ProLong^®^ Gold Antifade Mountant with DAPI (P36941, Life Technologies) or HOECHST 33342 (62249, Thermo Fisher) where specified. Images were acquired with a Leica SP2 confocal microscope equipped with a CCD camera.

### In situ hybridization

Whole mount RNA *in situ* hybridization on zebrafish embryos was performed as previously described (Thisse *et al.*, 1993). It is worth mentioning that treated and control embryos were hybridized together. *stat3* probe was obtained by PCR amplification from embryos cDNA using *stat3*_probe-fw (5′-TGCCACCAACATCCTAGTGT-3′) and *stat3*_probe-rv (5′-GCTTGTTTGCACTTTTGACTGA-3′) primers. *mt_nd2* probe was obtained by PCR amplification from embryos cDNA using *mt_nd2-fw* (5′-GCAGTAGAAGCCACCACAAA-3′) and *mt_nd2-rv* (5′-GGAATGCCGCGGATGTTATA-3′) primers. *pcna* probe was obtained as described by Baumgart *et al.* (2014). *sox9b* probe was obtained as described by Chiang *et al.* (2001). *her5* probe was obtained as described by Bally-Cuif, *et al.* (2000). *her4* probe was obtained as described by Takke *et al.* (1999). Fluorescence *in situ* hybridization was performed with FastBlue or TSA-amplification kit (Invitrogen) as described by Lauter *et al.* (2011).

### Transmission Electron Microscopy analysis

Larvae were anesthetised and fixed with 2.5% glutaraldehyde in 0.1 M sodium cacodylate buffer. After that, samples were dehydrated, embedded in epoxy resin, and prepared according to standard protocols by the Trasmission Electron Microscopy facility at the Department of Biology (University of Padova).

### Statistical analysis

Statistical analysis was performed with Graph Pad Prism V6.0. Data are presented as the means ± SEM and statistical analysis was determined by unpaired two tailed Student’s t-test. The p-values are indicated with the following symbols: *, p<0.05; **, p<0.01; ***, p < 0.001; ****, p<0.0001. For quantitative analysis, the sample size for each experiment was calculated assuming a Confidence Level of 95% (z-score 1,.96), a standard deviation of 0.5 and a Confidence Interval (margin of error) of 5%.

## Acknowledgments

We would like to thank Dr Luigi Pivotti, Dr Martina Milanetto, Dr Carlo Zatti, Dr Ludovico Scenna, and Mrs Shkendy Iljazi for their professional help in managing the Padua Zebrafish Facility, and Dr Andrea Vettori for his technical support. We are also grateful to Dr Valeria Poli and Dr Annalisa Camporeale for their kind gift of murine STAT3-encoding plasmids and their criticisms.

The work was fully supported by the AIRC grant IG 2017 19928 (to F.A.), with the contribution of the Telethon grant GGP19287 (to F.A.).

## Authors contribution

MP:acquisition, analysis, interpretation of data, drafting the work.

GM: acquisition, analysis, interpretation of data.

AD: acquisition, analysis, interpretation of data, drafting the work, critical revision.

LM: acquisition, analysis.

RMB: acquisition, analysis, interpretation of data, critical revision.

NF: acquisition, analysis, interpretation of data.

AT: acquisition, analysis.

NT: interpretation of data, drafting the work, critical revision.

GM: interpretation of data, drafting the work, critical revision.

FA: analysis, interpretation of data, drafting the work, critical revision.

---

**Fig. S1:**
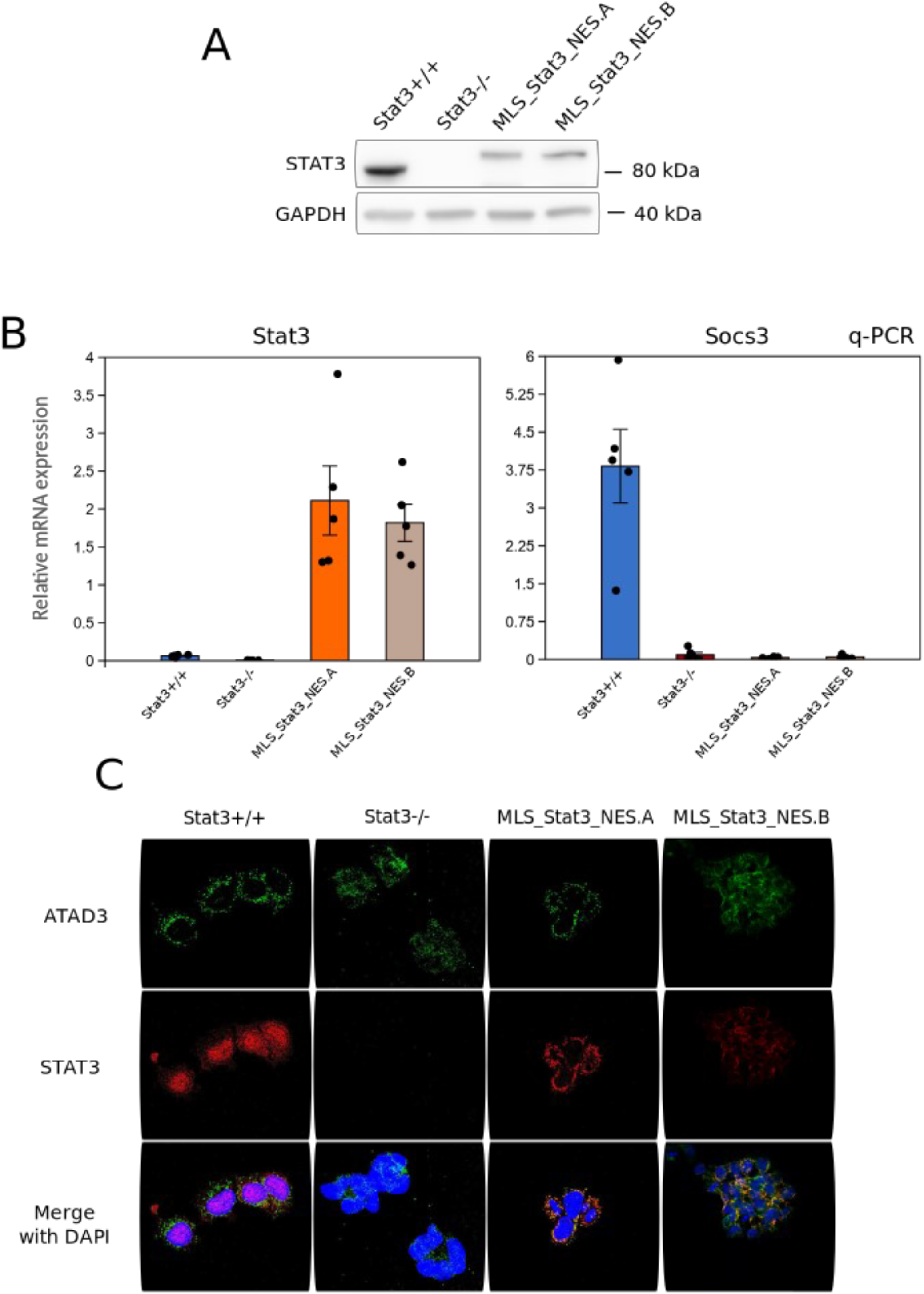
Validation of the *MLS_Stat3_NES* construct in murine Embryonic Stem Cells. **A:** Western blot for total STAT3 on *Stat3^+/+^, Stat3^-/-^* and MLS_Stat3_NES cells. Note the shift in molecular weight due to the presence of MLS and NES tags. STAT3 protein level in both MLS_Stat3_NES clones is lower than *Stat3^+/+^* cells. **B:** qPCR analysis of the Stat3 and its nuclear target gene *Socs3.* Gene expression analysis of *Stat3^+/+^* cells, *Stat3^-/-^* cells, and two MLS_Stat3_NES clones (A/B) cultured in presence of LIF. Note that both clones have the same undetectable level of *Socs3* as *Stat3^-/-^* cells. **C:** Representative confocal images of *Stat3^+/+^, Stat3^-/-^* and MLS_Stat3_NES cells stained with anti-STAT3 and anti-ATAD3 antibodies. Merge image shows co-localization between STAT3 and the nucleoids marked by ATAD3; DAPI serves as a nuclear counterstain. Scale bar: 20 μm.

**Fig S2:**
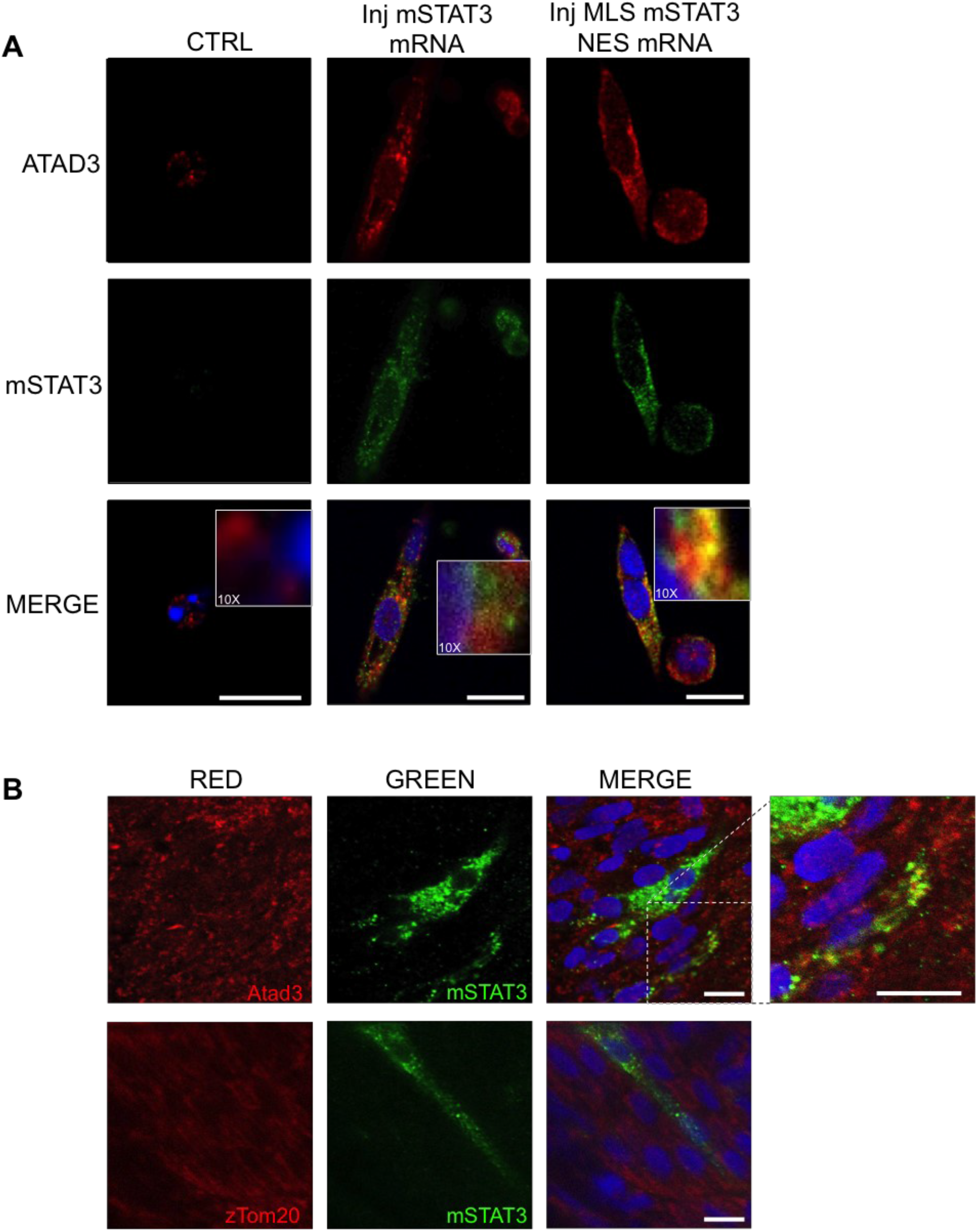
Validation of the injected mRNAs on zebrafish. **A:** IF on zebrafish cells, dissociated and plated from 24-hpf embryos injected with *mStat3* and *MLS_mStat3* mRNA. The antibody reveals the expression of mSTAT3 (green). The mito-targeted STAT3 co-localizes with ATAD3 (red), a marker of mitochondrial nucleoids, confirming the correct subcellular localization of the proteins. Conversely the analysis of cells from embryos injected with *mStat3* mRNA results in a more diffused staining. Scale bar = 10um. **B:** Whole mount IF on 24-hpf zebrafish embryos injected with pCS2 + MLS_mSTAT3_NES plasmid. The mosaic expression is driven by a CMV promoter to verify the intracellular localization of the murine protein. mSTAT3 (green) staining confirms the expected mitochondrial localization of the protein.

**Fig S3:**
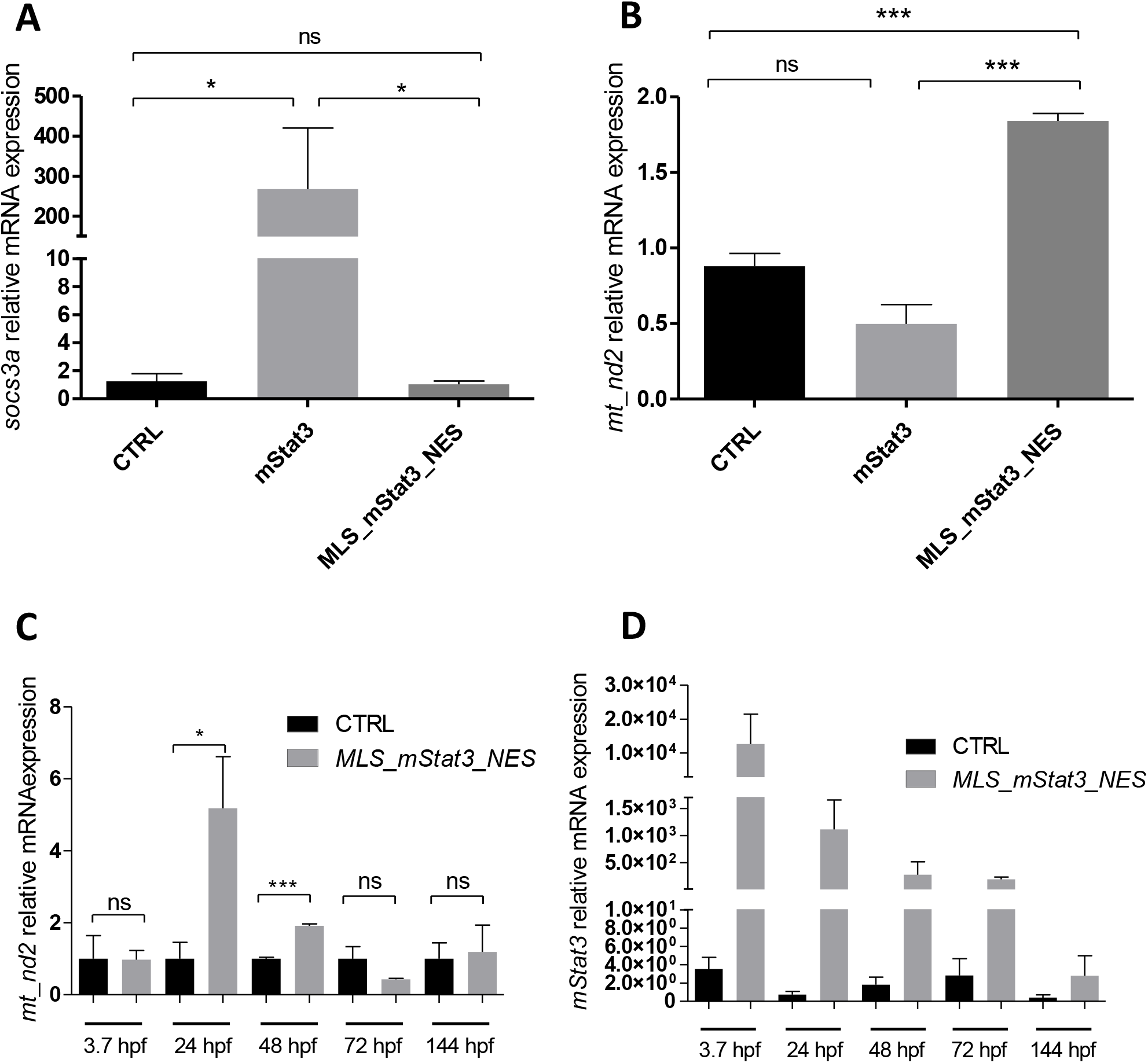
Validation of effects of *mStat3* and *MLS_Stat3_NES* mRNA injected in zebrafish embryos. **A:** qRT-PCR analysis of *socs3a* mRNA levels in 48-hpf embryos injected with *mStat3* and *MLS_mStat3_NES.* **B:** qRT-PCR analysis of *mt_nd2* mRNA levels in *mStat3* and *MLS_mStat3_NES* 48-hpf injected embryos. **C:** qRT-PCR analysis of *mt_nd2* levels from 3.7 hpf to 6 dpf, in larvae injected with *mStat3* mRNA. **D:** qRT-PCR analysis of *mStat3* levels from 3.7 hpf to 6 dpf, in larvae injected with *mStat3* mRNA. Statistical analysis was performed by unpaired t-test on 3 independent biological samples (where n not specified). *p<0.05; ***p<0.001; error bars=SEM.

**Fig. S4:**
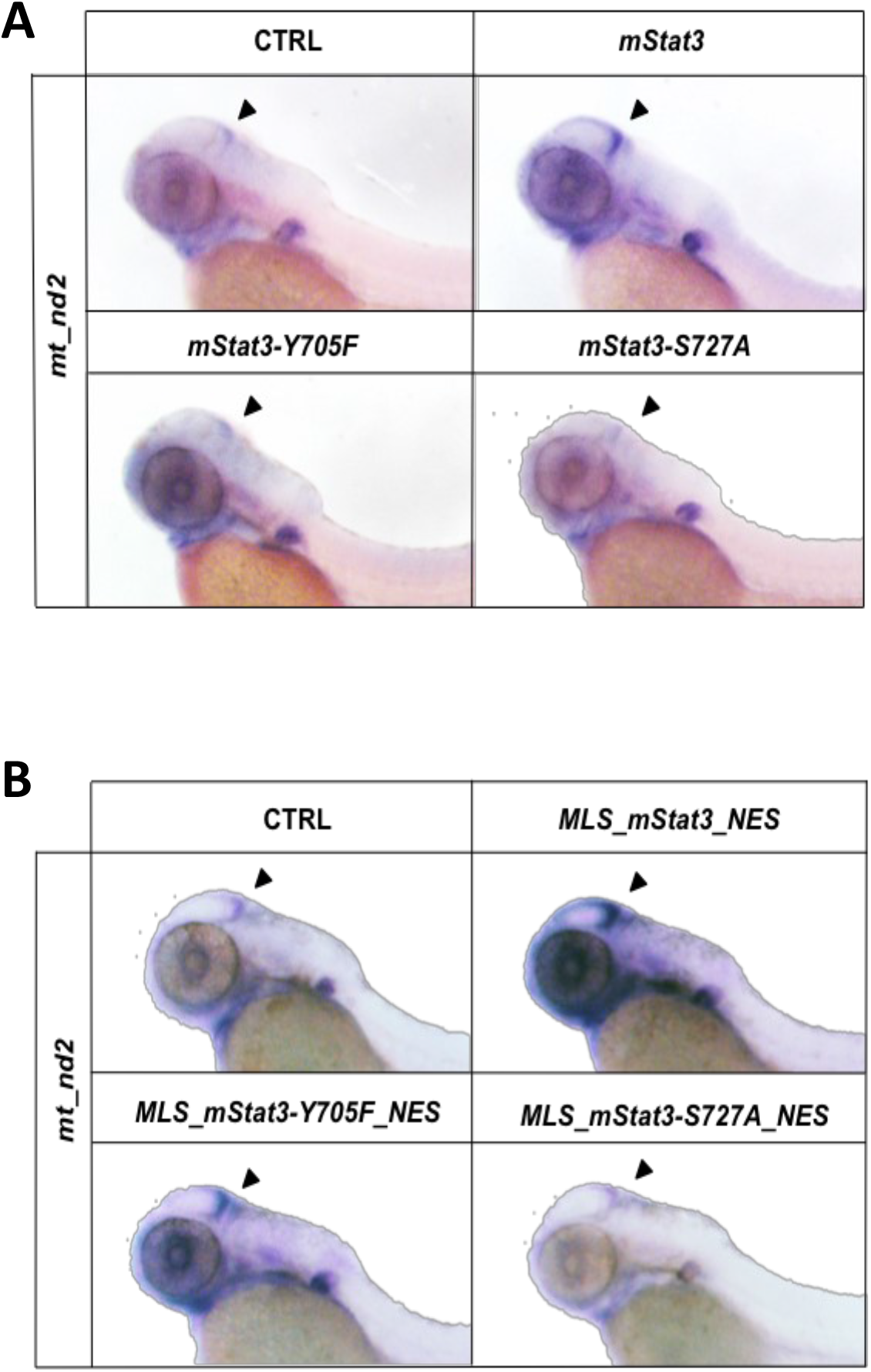
STAT3-dependent mitochondrial transcription depends on Y705 and S727 phosphorylations. **A:** WISH with *anti-mt_nd2* mRNA probe in 48-hpf uninjected embryos and embryos injected with either mStat3, mStat3-Y705F or mStat3-S727A. **B:** WISH with anti-*mt_nd2* mRNA probe in 48-hpf uninjected embryos and embryos injected with either MLS_mStat3_NES, MLS_mStat3_NES Y705F or MLS_mStat3_NES S727A.

**Fig. S5:**
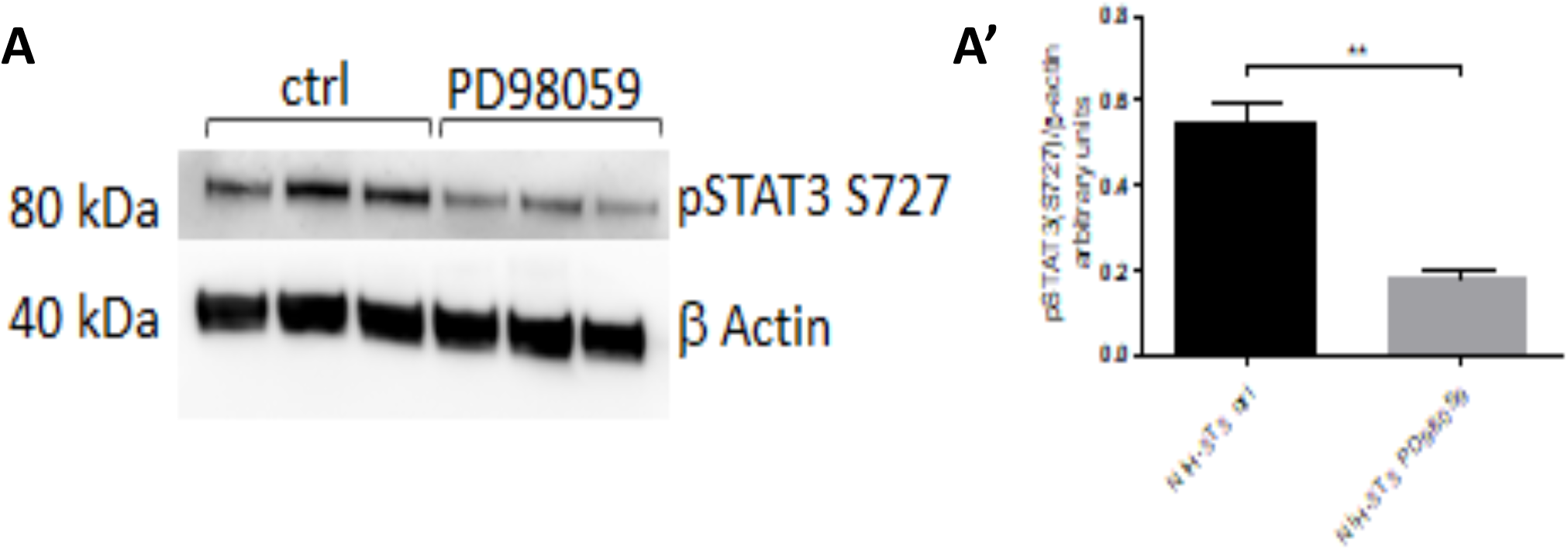
PD98059 inhibits S727 phosphorylation of STAT3 in NIH-3T3 cells. **A-A’:** western blot analysis of pSTAT3 S727 in NIH-3T3 cells treated for 24 hours with 12.5 μM PD98059 (ß-Actin was used as a loading control). Statistical analysis was performed by unpaired t-test on 3 independent biological samples. *p<0.05; error bars=SEM.

**Fig. S6:**
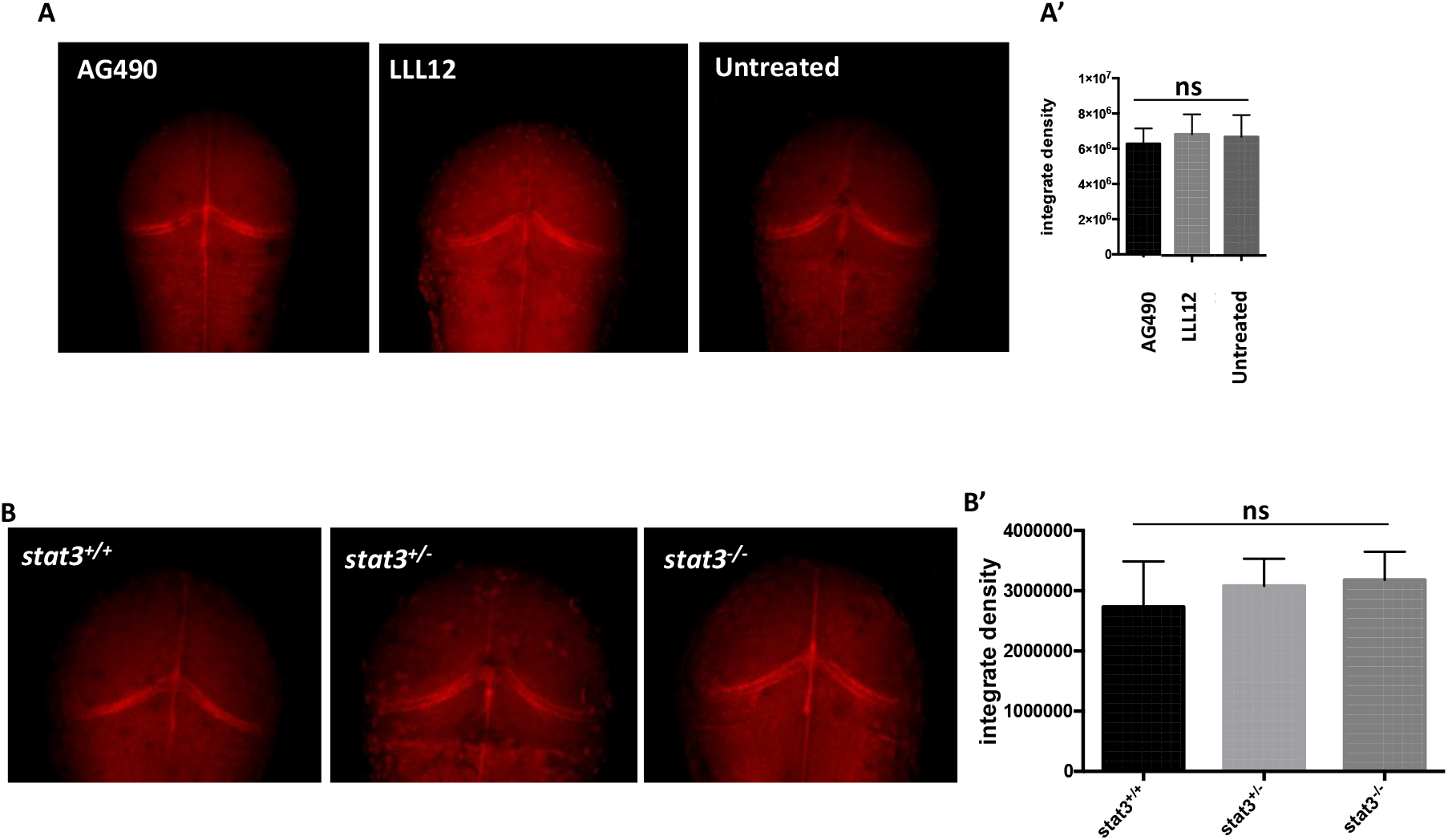
*mt_nd2* mRNA expression is not affected by AG-490 nor in 48-hpf *stat3* mutant larvae. **A-A’:** FISH with *mt_nd2* probe in the TeO of 48-hpf larvae treated for 24 hours with AG490. Fluorescence quantification of *mt_nd2* mRNA levels in the TeO (n=10). **B:** FISH with *mt_nd2* probe in the TeO of 48-hpf *stat3^+/+^, stat3^+/-^*, and *stat3^-/-^* larvae. B’: fluorescence quantification of *mt_nd2* mRNA levels in the TeO. Statistical analysis was performed by unpaired t-test on 3 independent biological samples. ns = not significant; error bars=SEM.

**Fig. S7:**
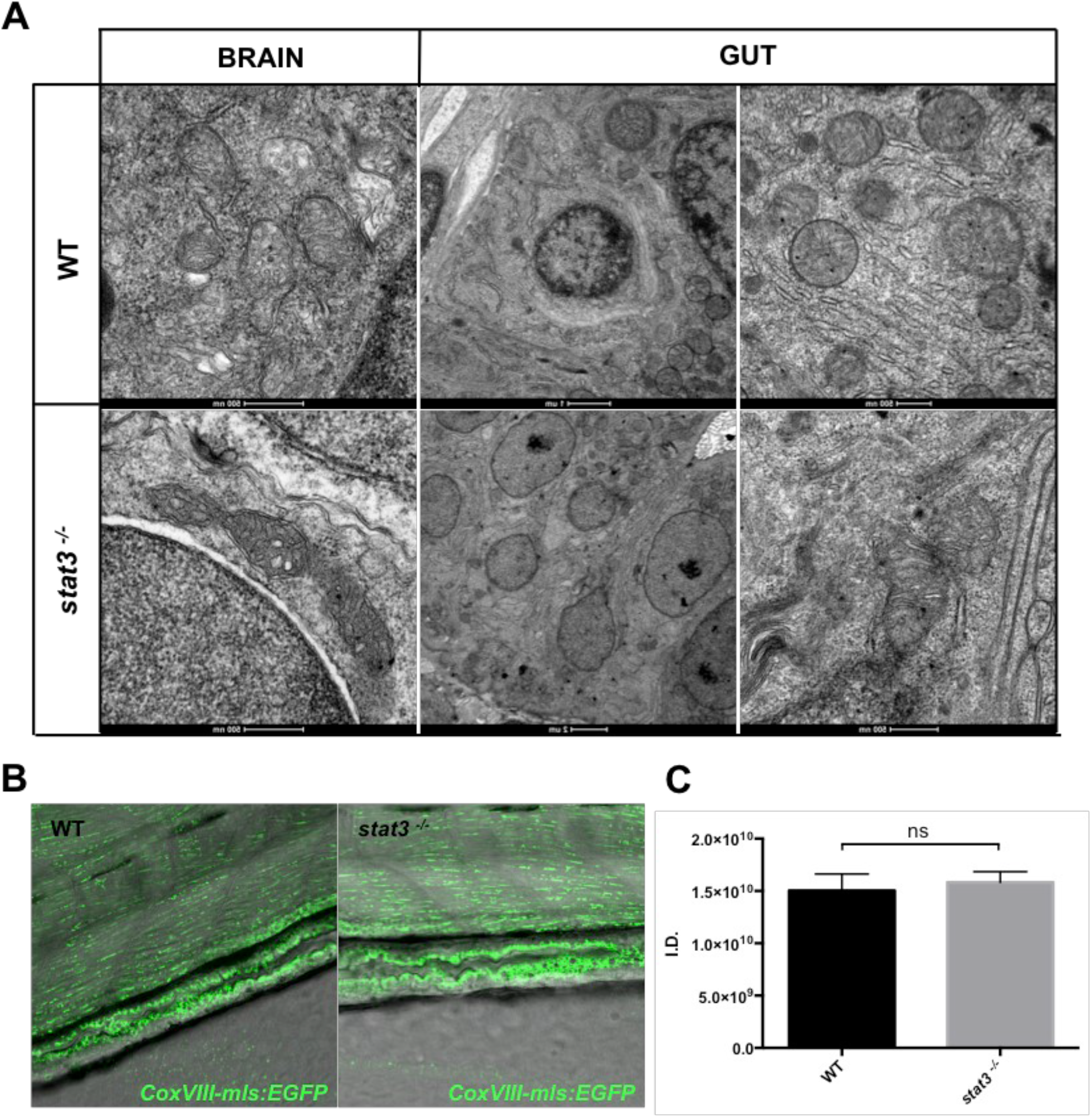
Stat3 depletion does not affect mitochondria morphology and biogenesis in the brain and intestine of *stat3^-/-^* larvae. **A:** TEM analysis of mitochondrial morphology in intestine and brain of 6-dpf *stat3^-/-^* mutants and WT siblings. **B:** EGFP expression in the intestine of 6-dpf *stat3^-/-^/Tg(CoxVIII-mls:EGFP)* and *WT/Tg(CoxVIII-mls:EGFP)* siblings (n=6). **C:** Fluorescence quantification of EGFP expression in the intestine of 6-dpf *stat3^-/-^/Tg(CoxVIII-mls:EGFP)* and *WT/Tg(CoxVIII-Emls:EGFP)* siblings (p-value= 0.,6878). Statistical analysis was performed by unpaired t-test on indicated number of samples; ns = not significant; error bars=SEM.

## REFERENCES

Alessi D.R., Cuenda A., Cohen P., Dudley D.T., Saltiel A.R. (1995). PD 098059 is a specific inhibitor of the activation of mitogen-activated protein kinase kinase in vitro and in vivo. J Biol Chem. 270(46):27489–94.

Antico Arciuch V.G., Elguero M.E., Poderoso J.J., Carreras M.C. (2012). Mitochondrial regulation of cell cycle and proliferation. Antioxid Redox Signal. 16(10: 1150–80.

Avalle L., Camporeale A., Morciano G., Caroccia N., Ghetti E., Orecchia V., Viavattene D., Giorgi C., Pinton P., Poli V. (2019). STAT3 localizes to the ER, acting as a gatekeeper for ER-mitochondrion Ca2+ fluxes and apoptotic responses. Cell Death Differ. May;26(5):932–942.

Bally-Cuif, L., Goutel, C., Wassef, M., Wurst, W., & Rosa, F. (2000). Coregulation of anterior and posterior mesendodermal development by a hairy-related transcriptional repressor. Genes & development. 14(13), 1664–77.

Baumgart, M., Groth, M., Priebe, S., Savino, A., Testa, G., Dix, A., Ripa, R., Spallotta, F., Gaetano, C., Ori, M., Terzibasi Tozzini, E., Guthke, R., Platzer, M. and Cellerino, A. (2014). RNA-seq of the aging brain in the short-lived fish N. furzeri - conserved pathways and novel genes associated with neurogenesis. Aging Cell. 13: 965–974.

Becker S., Groner B., Müller C.W. (1998). Three-dimensional structure of the Stat3beta homodimer bound to DNA. Nature. 394:145–151 10.1038/28101

Burdon T., Chambers I., Stracey C., Niwa H., Smith A. (1999). Signaling mechanisms regulating self-renewal and differentiation of pluripotent embryonic stem cells. Cells Tissues Organs. 165(3-4):131–43.

Carbognin, E., Betto, R. M., Soriano, M. E., Smith, A. G. and Martello, G. (2016). Stat3 Promotes Mitochondrial Transcription and Oxidative Respiration during Maintenance and Induction of Naive Pluripotency. EMBO J. 35, 618–634.

Chiang E.F.L., Pai C.I., Wyatt M., Yan Y.L., Postlethwait J., and Chung B.C. (2001). Two sox9 genes on duplicated zebrafish chromosomes: Expression of similar transcription activators in distinct sites. Dev Bio. 231: 149–163.

Decker T., & Kovarik P. (2000). Serine phosphorylation of STATs. Oncogene. 19: 2628–2637.

Facchinello N., Skobo T., Meneghetti G., Colletti E., Dinarello A., Tiso N., Costa R., Gioacchini G., Carnevali O., Argenton F., Dalla Valle L. (2017) *nr3c1* null mutant zebrafish are viable and reveal DNA-binding-independent activities of the glucocorticoid receptor. Scientific reports. 7: 4371.

Feng J.Y., Xu Y., Barauskas O., Perry J.K., Ahmadyar S., Stepan G., Yu H., Babusis D., Park Y., McCutcheon K., Perron M., Schultz B.E., Sakowicz R., Ray A.S. (2015) Role of Mitochondrial RNA Polymerase in the Toxicity of Nucleotide Inhibitors of Hepatitis C Virus. Antimicrob Agents Chemother. 60(2):806–17.

Fouse S.D., Costello J.F. (2013). Cancer Stem Cells Activate STAT3 the EZ Way. Cancer Cell. 23(6). 711–713.

Gagnon J.A., Valen E., Thyme S.B., Huang P., Ahkmetova L., Pauli A., et al. (2014). Efficient Mutagenesis by Cas9 Protein-Mediated Oligonucleotide Insertion and Large-Scale Assessment of Single-Guide RNAs. PLoS ONE. 9(5): e98186.

Ghoshal S., Fuchs B. C., & Tanabe K. K. (2016). STAT3 is a key transcriptional regulator of cancer stem cell marker CD133 in HCC. Hepatobiliary Surgery and Nutrition. 5(3), 201–203.

Gough D.J., Corlett A., Schlessinger K., Wegrzyn J., Larner A.C., Levy DE. (2009). Mitochondrial STAT3 supports Ras-dependent oncogenic transformation. Science. 324(5935:1713–6.

Gough D.J., Koetz L., Levy D.E. (2013) The MEK-ERK Pathway Is Necessary for Serine Phosphorylation of Mitochondrial STAT3 and Ras-MediatedTransformation. PLoS ONE 8(11: e83395.

Grivennikov, S., Karin, E., Terzic, J., Mucida, D., Yu, G. Y., Vallabhapurapu, S., Scheller, J., Rose-John, S., Cheroutre, H., Eckmann, L. et al. (2009). IL-6 and Stat3 are Required for Survival of Intestinal Epithelial Cells and Development of Colitis-Associated Cancer. Cancer. Cell. 15, 103–113.

Gurbuz V., Konac E., Varol N., Yilmaz A., Gurocak S., Menevse S., Sozen S. (2014). Effects of AG490 and S3I-201 on regulation of the JAK/STAT3 signaling pathway in relation to angiogenesis in TRAIL-resistant prostate cancer cells in vitro. Oncol Lett. 7(3): 755–763.

Horvath C.M., Wen Z., Darnell J.E., Jr. (1995). A STAT protein domain that determines DNA sequence recognition suggests a novel DNA-binding domain. Genes Dev. 9:984–994.

Huang G., Yan H., Ye S., Tong C., Ying Q.L. (2014). STAT3 phosphorylation at tyrosine 705 and serine 727 differentially regulates mouse ESC fates. Stem Cells. 32:1149–1160.

Johnston P.A., Grandis J.R. (2011). STAT3 signaling: anticancer strategies and challenges. Mol Interv. 11(1): 18–26.

Kimmel C.B., Ballard W.W., Kimmel S.R., Ullmann B., Schilling, T.F. (1995), Stages of embryonic development of the zebrafish. Dev. Dyn. 203: 253–310.

Lauter G., Söll I., Hauptmann G. (2011). Two-color fluorescent in situ hybridization in the embryonic zebrafish brain using differential detection systems. BMC developmental biology. 11, 43.

Liu Y., Sepich D.S., Solnica-Krezel L. (2017) Stat3/Cdc25a-dependent cell proliferation promotes embryonic axis extension during zebrafish gastrulation. PLoS Genet. 13(2): e1006564. https://doi.org/10.1371/journal.pgen.1006564

Livak K.J., & Schmittgen B.M. (2001) Analysis of relative gene expression data using real-time quantitative PCR and the 2(-Delta Delta C(T)) method. Methods. 25(4): 402–8.

Macias E., Rao D., Carbajal S., Kiguchi K. and DiGiovanni, J. (2014). Stat3 Binds to mtDNA and Regulates Mitochondrial Gene Expression in Keratinocytes. J. Invest. Dermatol. 134, 1971–1980.

Mantel C., Messina-Graham S., Moh A., Cooper S., Hangoc G., Fu X.Y., Broxmeyer H. E. (2012). Mouse hematopoietic cell-targeted STAT3 deletion: stem/progenitor cell defects, mitochondrial dysfunction, ROS overproduction, and a rapid aging-like phenotype. Blood. 120(13), 2589–99.

Martello G., Bertone P., Smith A. (2013) Identification of the missing pluripotency mediator downstream of leukaemia inhibitory factor. EMBO J. 32(19:2561–74.

Martorano L., Peron M., Laquatra C., Lidron E., Facchinello N., Meneghetti G., Tiso N., Rasola A., Ghezzi D., Argenton F. (2019) The zebrafish orthologue of the human hepatocerebral disease gene *MPV17* plays pleiotropic roles in mitochondria. Dis Model Mech. 12(3): dmm0372226.

Matsuda T., Nakamura T., Nakao K., Arai T., Katsuki M., Heike T., Yokota T. (1999). STAT3 activation is sufficient to maintain an undifferentiated state of mouse embryonic stem cells. EMBO J. 18(15):4261–9.

Matthews J.R., Sansom O.J. and Clarke A.R. (2011). Absolute Requirement for STAT3 Function in Small-Intestine Crypt Stem Cell Survival. Cell Death Differ. 18, 1934–1943.

Meier J.A., Larner A.C. (2014). Toward a new STATe: the role of STATs in mitochondrial function. Seminars in immunology, 26(1), 20–8.

Mohr A., Fahrenkamp D., Rinis N., Muller-Newen G. (2013) Dominant-negative activity of the STAT3-Y705F mutant depends on the N-terminal domain. Cell Commun Signal. 11: 83.

Ni C., Hsieh H., Chao Y., Wang D.L. (2004). Interleukin-6-induced JAK2/STAT3 signaling pathway in endothelial cells is suppressed by hemodynamic flow. American Journal of Physiology. Cell Physiology. 287(3) C771–C780.

Oates A.C., Wollberg P., Pratt S.J., Paw B.H., Johnson S.L., Ho R.K., Postlrthwait J.H., Zon L.I., Wilks A.F. (1999). Zebrafish stat3 is expressed in restricted tissues during embryogenesis and stat1 rescues cytokine signaling in a STAT1-deficient human cell line. Dev Dyn. 215:352–370.

O’Shea J.J., Schwartz D.M., Villarino A.V., Gadina M., McInnes I.B., Laurence A. (2015). The JAK-STAT pathway: impact on human disease and therapeutic intervention. Annu Rev Med. 66:311–28.

Park J.S., Lee J., Lim M.A., Kim E.K., Kim S.M., Ryu J.G., Lee J.H., Kwok S.K., Park K.S., Kim H.Y., Park S.H., Cho M.L. (2014) JAK2-STAT3 blockade by AG490 suppresses autoimmune arthritis in mice via reciprocal regulation of regulatory T Cells and Th17 cells. J Immunol. May 1;192(9):4417–24.

Peron M., Dinarello A., Meneghetti G., Martorano L., Facchinello N., Vettori A., Licciardello G., Tiso N., Argenton F. (2020) The stem-like STAT3-responsive cells of zebrafish intestine are WNT/β-catenin dependent. Development. 147(12): dev188987.

Qin H.R., Kim H.J., Kim J.Y., Hurt E.M., Klarmann G.J., Kawasaki B.T., Duhagon Serrat M.A., Farrar W.L. (2008). Activation of signal transducer and activator of transcription 3 through a phosphomimetic serine 727 promotes prostate tumorigenesis independent of tyrosine 705 phosphorylation. Cancer research. 68(19), 7736–41.

Shi X., Zhang H., Paddon H., Lee G., Cao X., Pelech S. (2006). Phosphorylation of STAT3 serine-727 by cyclin-dependent kinase 1 is critical for nocodazole-induced mitotic arrest. Biochemistry. 45:5857–5867.

Smith A.G., Heat J.K., Donaldson D.D., Wong G.G., Moreau J., Stahl M., Rogers D. (1988) Inhibition of pluripotential embryonic stem cell differentiation by purified polypeptides. Nature. 336(6200): 688–690.

Strutt S.C., Torrez R.M., Kaya E., Negrete O.A., Doudna J.A. (2018) RNA-dependent RNA targeting by CRISPR-Cas9. eLife. 7: e32724.

Szczepanek K., Chen, Q., Derecka M., Salloum F.N., Zhang Q., Szelag M., Cichy J., Kukreja R.C., Dulak J., Lesnefsky E.J., et al. (2011). Mitochondrial-targeted signal transducer and activator of transcription 3 (stat3) protects against ischemia-induced changes in the electron transport chain and the generation of reactive oxygen species. J. Biol. Chem. 286, 29610–29620.

Taanman, J. (1999). The mitochondrial genome: structure, transcription, translation and replication. Biochimica et Biophysica Acta-Bioenergetics. 1410(2), 103–123.

Takke C., Dornseifer P., v Weizsäcker E., Campos-Ortega J.A. (1999). her4, A zebrafish homologue of the Drosophila neurogenic gene E(spl), is a target of NOTCH signalling. Development. 126: 1811–1821.

Thisse B., Pflumio S., Fürthauer M., Loppin B., Heyer V., Degrave A., Woehl R., Lux A., Steffan T., Charbonnier X.Q. and Thisse C. (2001) Expression of the zebrafish genome during embryogenesis (NIH R01 RR15402). ZFIN Direct Data Submission. (http://zfin.org)

Tian Z.J., An W. (2004) ERK1/2 contributes negative regulation to STAT3 activity in HSS-transfected HepG2 cells. Cell Res. Apr;14(2):141–7.

Wang J., Zhou M., Jin X., Li B., Wang C., Zhang Q., Liao M., Hu X., Yang M. (2019) Glycochenodeoxycholate induces cell survival and chemoresistance via phosphorylation of STAT3 at Ser727 site in HCC. J Cell Physiol. 2019;1–12.

Wegrzyn J., Potla R., Chwae Y.J., Sepuri N.B., Zhang Q., Koeck T., Derecka M., Szczepanek K., Szelag M., Gornicka A., Moh A., Moghaddas S., Chen Q., Bobbili S., Cichy J., Dulak J., Baker D.P., Wolfman A., Stuehr D., Hassan M.O., Fu X.Y., Avadhani N., Drake J.I., Fawcett P., Lesnefsky E.J., Larner A.C. (2009). Function of mitochondrial Stat3 in cellular respiration. Science. 323(5915):793–7.

Wei W., Tweardy D.J., Zhang M., Zhang X., Landua J., Petrovic I., Bu W., Roarty K., Hilsenbeck S.G., Rosen J.M., Lewis M.T. (2014). STAT3 signaling is activated preferentially in tumor-initiating cells in claudin-low models of human breast cancer. Stem Cells. 32: 2571–2582.

